# *Helicobacter* small RNA regulates host adaptation and carcinogenesis

**DOI:** 10.1101/2020.02.15.950279

**Authors:** Ryo Kinoshita-Daitoku, Kotaro Kiga, Ryota Otsubo, Yoshitoshi Ogura, Takahito Sanada, Zhu Bo, Tuan Vo Phuoc, Tokuju Okano, Tamako Iida, Rui Yokomori, Eisuke Kuroda, Sayaka Hirukawa, Mototsugu Tanaka, Arpana Sood, Phawinee Subsomwong, Hiroshi Ashida, Tran Thanh Binh, Lam Tung Nguyen, Khien Vu Van, Dang Quy Dung Ho, Kenta Nakai, Toshihiko Suzuki, Yoshio Yamaoka, Tetsuya Hayashi, Hitomi Mimuro

## Abstract

Type-1 carcinogenic *Helicobacter pylori* that is known to evolve during long-term infection, enters the stomach orally and causes gastric cancer using the carcinogenic protein CagA^1^. However, little is known about the adaptation mechanisms of *H. pylori* when the environment changes from the outside to the inside of the living body. Here we show that small non-coding RNA HPnc4160 is a crucial novel RNA molecule of *H. pylori* that negatively regulates bacterial-host adaptation and gastric cancer. *H. pylori* isolated from gerbil’s stomachs eight weeks post-infection acquired mutations in the increased number of T-repeats within the upstream region of the HPnc4160 coding region, which leads to reduced HPnc4160 expression levels that also seen in cancer patients-derived *H. pylori*. By comparing RNA-seq and iTRAQ analysis between wild-type and *hpnc4160* deficient mutant strains, we identified eight targets of HPnc4160 including *cagA* and unknown factors. Mice infection experiment revealed that the *hpnc4160* deficient mutant had a higher number of colonized bacteria in the mice stomach than the wild-type strain, indicating that reduced expression levels of HPnc4160 was important for bacterial host adaptation. The expression level of HPnc4160 was lower in the clinical isolates derived from gastric cancer patients compared with non-cancer-derived strains, while the mRNA expression levels of target factors were higher. Our findings highlight the first discovery that HPnc4160 is an important small RNA for bacteria to adapt to the host environment leading to gastric carcinogenesis.

## Main

*Helicobacter pylori* infection has a very high prevalence that about half of the world population is infected. Patients infected with CagA-positive *H. pylori* closely related to disease malignancy have been reported to have an increased risk of peptic ulcer, chronic gastritis, intestinal metaplasia, and gastric cancer.^2, 3^ The highly pathogenic *H. pylori* possess a *cag* pathogenicity island that encodes components of a Type IV secretion system (TFSS), which is an injection needle, and a carcinogenic factor CagA effector protein. The *H. pylori* that has reached the gastric epithelium injects CagA protein, peptidoglycan, and heptose-1,7-bisphosphate into the attached cells via TFSS, and stimulates signal transduction pathways such as NF-κB to promote production of chemokines such as interleukin-8 (IL-8)^4–11^.

It is considered that the genetic diversity at the genomic level that is characteristic of *H. pylori* is very important to establish a persistent infection in an infected host with different backgrounds, adapting to a gastric niche with severe environmental changes^12^. The *H. pylori* gene mutations are characterized by the presence of simple repetitive sequences such as mononucleotide repeats (such as poly-T) and dinucleotide repeats (such as CT-repeats). From the analysis of clinical isolates of *H. pylori* so far, it is considered that phase variations are induced by ON / OFF control of gene expression of such as outer membrane proteins (OMPs) due to expansion and contraction of these simple repetitive sequences^12–14^. In the course of chronic infection, in order to escape from the host immunity, it is assumed that strong diversity in *H. pylori* OMPs that can serve as highly antigenic cell surface antigens will cause a strong directivity of selection^15^. Therefore, to understand the mechanism of persistent infection by *H. pylori*, we analyzed bacterial gene mutations acquired by *H. pylori* in the course of persistent infection using an experimental animal infection system with the same host genetic background. With growing evidence that bacterial small RNA (sRNA)-mediated target gene expression in response to changes in the environment, we focused sRNAs as well^16^.

## Identification of HPnc4160

To analyze the bacterial gene mutation acquired by *H. pylori* during the persistent infection, Mongolian gerbils (n=10) were inoculated with *H. pylori* ATCC 43504 wild-type strain for 8 wks. *H. pylori* in the infected stomachs were isolated (n=40; Fig. 1a), and analyzed comparative whole genome sequences (Fig. 1b, Supplementary Information 1, 2). We totaled genomic positions, where these mutations were introduced, for each coding and intergenic region, and identified 13 regions (Regions R1, R3-R5, R7-R8, R10-R16) where mutations were introduced in 50% or more of the strains (Fig. 1b, Extended Data Table 1).

**Fig. 1:**
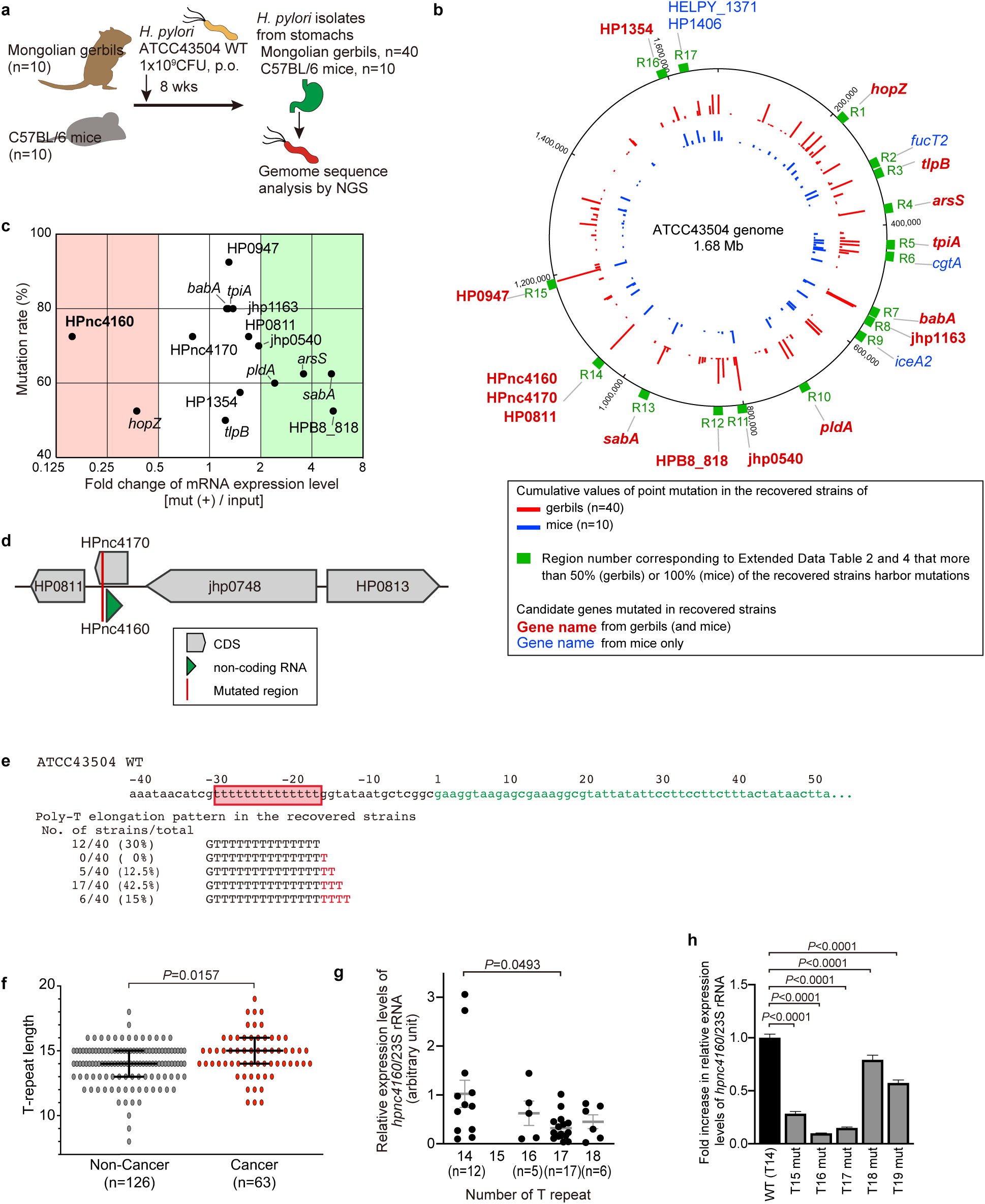
*H. pylori* acquire poly-T extension in upstream of HPnc4160 small RNA to decrease its expression during infection *in vivo*. **a**, Experimental strategies schematic. **b**, Circular genomic map of ATCC 43504 strain recovered from stomachs of gerbils and mice. **c**, Mutation rates and expression levels of candidate RNAs (mRNA or non-coding RNA). RNA expression levels of the ORFs or nearby genes of genome regions [total 15 genes (Extended Data Fig. 1a), which mutated in more than 50% of the recovered strains from the gerbils, were assessed and plotted against the mutation rates. **d**, Schematic structures of genes around HPnc4160. **e**, Schematic diagram of genome DNA sequence around the HPnc4160 and poly-T sequence of recovered strains harboring mutations. Green indicates HPnc4160 transcribed sequence, red frame indicates poly-T mutated stretches in the upstream region of HPnc4160 of ATCC 43504 wild-type (WT) strain, and red-colored “T” indicated inserted nucleotides of each recovered strain compared with WT. **f**, The T-repeat length in the upstream region of HPnc4160 of clinical isolates. Isolated strains from cancer patients (Cancer) have a higher number of T-repeat in the upstream region of Hpnc4160 compared with the isolates from non-cancer patients (Non-Cancer). Data are presented as means with 95% confidence interval. *P* values represent the results of the two-tailed Mann-Whitney test. **g**, Expression levels of HPnc4160 in the recovered *H. pylori* strains from Mongolian gerbils (n=40). Relative expression levels of HPnc4160 were measured by real-time PCR and plotted against T-repeat length in the upstream region of HPnc4160. Data are presented as means with s.d. *P* values represent the results of two-tailed Dunn’s multiple comparison test. **h**, The relative expression levels of HPnc4160 in the *H. pylori* strains genetically modified with the T-repeat length. Data are presented as means with s.d. (n=3). *P* values are from Dunnett’s multiple comparison test (at two-sided).

To investigate whether the mutated region affects RNA expression in isolates recovered from the gerbils, the expression levels of mRNA or non-coding RNA around the mutated regions were quantified by quantitative PCR. Among the corresponding 15 CDSs and non-coding RNA (HP0947, *babA*, *tpiA*, jhp1163, HP0811, HPnc4160, HPnc4170, jhp0540, *araS*, *pldA*, *sabA*, HP1354, *hopZ*, *tlpB*, HPB8_818), we found that the expression level of HPnc4160, a non-coding small RNA (sRNA) of unknown function, fluctuated the most compared to the wild-type (Fig. 1c, Extended Data Fig. 1a). Similar results were also obtained with the strains (n=10) isolated from C57BL/6 mice (n=5) (Fig. 1b, Extended Data Fig. 1a, b; Supplementary Information 3, 4).

Region R14 is the upstream region of HPnc4160 and HP0811, and is located in the CDS of HPnc4170 (*aapB* small ORF homologue) encoded by the complementary sequence of HPnc4160 (Fig. 1d) ^17^. HPnc4160 and its upstream T repeat were conserved in various *H. pylori* genome analysis strains, and T repeat length was different depending on the strain (Extended Data Fig. 2a). A repeat of 2 to 4 bases of thymidine was inserted into the repeat of isolates from rodents, and the repeat length increased depending on the infection period (Fig. 1e, Extended Data Fig. 2b-e). Importantly, sequence analysis of clinical isolates showed that T repeat lengths were longer in cancer patient-derived strains than in non-cancer patient-derived strains (Fig. 1f, Supplementary Information 5). However, expansion of the repeat was not observed in long-term *in vitro* culture (Extended Data Fig. 3). Next, we analyzed the change in HPnc4160 sRNA expression levels by T repeat length. In strains recovered from *H. pylori-*infected rodent stomachs, HPnc4160 expression level tended to decrease with expansion of T repeat length (Fig. 1g). To exclude the effects of mutations other than the T repeats, we further analyzed RNA expression levels of HP0811, HPnc4170 and HPnc4160 in various mutants in which the T repeat sequence was inserted into the HPnc4160 upstream region of wild-type strain (T15mut, T16mut, T17mut, T18mut, and T19mut). In strains in which the number of T repeats was greater than T14 of wild-type, the expression levels of HPnc4160, but not HP0811 and HPnc4170, were significantly reduced compared with wild-type (Fig. 1h, Extended Data Fig. 4a-c,). Although HPnc4160 and HPnc4170 were initially reported as the small ORF-encoding mRNA/antisense RNA family aapB/IsoB, in which the Iso transcript acts as asRNA antitoxin to modify the aap expression^18^, our data indicated that HPnc4160 expression levels had no effect on HPnc4170 levels. These results indicated that expression levels of HPnc4160 sRNA decreased when the number of Region R14 T repeats increased due to persistent intragastric infection.

## Identification of HPnc4160 target genes

Many sRNAs regulate the expression of a target mRNA by specifically binding to a complementary sequence in the target protein coding mRNA. To elucidate the target mRNA of HPnc4160, we made a Δ*hpnc4160*/*hpnc4170* strain, in which both the HPnc4160 and the HPnc4170 on the complementary strand were deleted, and analyzed comparative mRNA and protein expression. We identified eight factors (*cagA*, *hofC*, HELPY_1262, *hpaA*, *horB*, *omp14*, *hopE*, and HP1227) with P-values lower than 0.001 (RNA-seq analysis) and 0.01 [isobaric tag for relative and absolute quantitation (iTRAQ) labeling and LC-MS/MS analysis] (Fig. 2a-c, Extended Data Table 2a-b). Of these, *cagA* was prominent in expression levels of mRNA and protein.

**Fig. 2:**
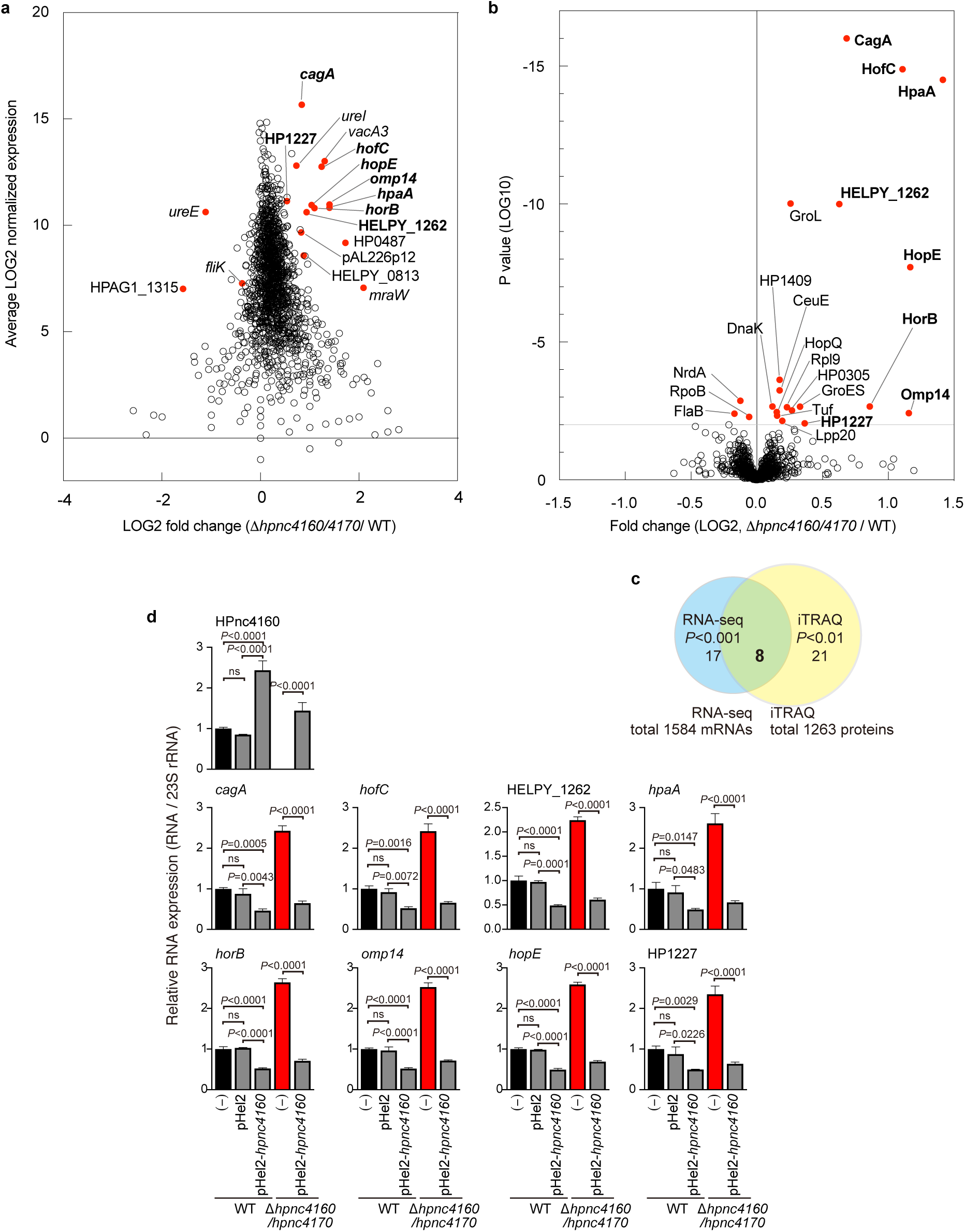
HPnc4160 downregulates expression levels of bacterial pathogenic factors. **a**, The MA plot of ratios [Δ*hpnc4160*/*hpnc4170* / wild-type (WT) *H. pylori*] versus their normalized average mRNA expression determined by RNA-sequencing (RNA-Seq). The red dots showed genes of *P*<0.001. **b**, Volcano plots of the proteins quantified by isobaric tags for relative and absolute quantification (iTRAQ) analysis comparing WT and Δ*hpnc4160*/*4170*. Each point represents the difference in expression (fold change) between the two groups plotted against the level of statistical significance. The red dots showed proteins of *P*<0.01. **c**, The Venn diagram represents the number of factors whose expression exhibited significant differences between Δ*hpnc4160*/*hpnc4170* and WT. **d**, The relative RNA expression levels of target candidates of HPnc4160 showed an inverse correlation with HPnc4160. The results represent the average of three separate experiments (each n=3). Data are presented as means ± s.d. (error bars). *P* values are from Tukey’s multiple comparison test (at two-sided). ns: not significant.

We analyzed whether the mRNA expression levels of the identified eight factors depend on the presence of HPnc4160. Although the expression level of HPnc4160 showed a decreasing trend with T16mut as the lowest value, the mRNA of the eight candidates showed increasing trends with T16mut as the highest value. The Spearman correlation coefficients r between the expression levels of these target mRNAs and HPnc4160 showed a very strong inverse correlation from −0.7714 to −1.0 (Extended Data Fig. 4d).

Next, we examined the mRNA expression correlation between HPnc4160 and each factor. The HPnc4160 overexpression strain (WT / pHel2-*hpnc4160*) significantly increased the expression level of HPnc4160 compared to the wild-type strain, but significantly decreased the expression level of each factor mRNA. On the other hand, in Δ*hpnc4160*/*hpnc4170* strain, mRNA expression of each target increased compared to the wild-type. Since Δ*hpnc4160*/*hpnc4170* strain also lacks the HPnc4170 sequence present in the complementary strand of HPnc4160, we constructed a Δ*hpnc4160*/*hpnc4170* /pHel2-*hpnc4160* strain complementing only HPnc4160 to confirm the effect of the HPnc4170 sequence on HPnc4160 target mRNA expression. Compared to the Δ*hpnc4160*/*hpnc4170* strain, the mRNA expression levels of the candidates were decreased in the HPnc4160 complemented Δ*hpnc4160*/*hpnc4170* /pHel2-*hpnc4160* (Fig. 2d, Extended Data Fig. 5a). These data indicated that the expression levels of the HPnc4160 target 8 candidates were suppressed depending on the expression level of HPnc4160.

When sRNA binds within a few bases around the 5 ′UTR or start codon of target RNA, it often competes with ribosomes and inhibits translation initiation. If the sRNA binds within the CDS far downstream of the initiation codon, it causes mRNA degradation by RNase E or RNase III to suppress translational activation^19^. We confirmed by electrophoretic mobility shift assays (EMSA) that HPnc4160 binds to the 5 ′UTR of seven genes other than *cagA* (Fig. 3a). In seven factors other than *cagA*, we confirmed a sequence complementary to the HPnc4160 sequence in the 5 ′UTR of each gene (Extended Data Fig. 5b-c). In the *cagA* gene, we found HPnc4160-binding sequences Type 1 at one position (2344 nt), and Type 2 at four positions (2838, 2940, 3042, and 3144 nt) within the CM/CRPIA motifs in CagA C-terminal region, which is known to bind with host signal proteins^7, 20^ (Fig. 3b, Extended Data Fig. 5d-f). We confirmed direct binding of *cagA* partial CDS (positions 2778 to 3236 nt from start codon of *cagA*) to HPnc4160 (Fig. 3c). The binding between the two was abolished in the NB-*cagA* RNA in which the HPnc4160 binding sequence was mutated at four positions (Type 2) while the amino acid sequence of CagA was preserved (Fig. 3c, Extended Data Fig. 5g). In addition, we found that in the presence of *H. pylori* RNase III recombinant protein, the binding between HPnc4160 and biotin-labeled partial *cagA* mRNA, but not NB-*cagA* RNA, disappeared (Fig. 3d, Extended Data Fig. 5i). These data clearly demonstrated that HPnc4160 controls *cagA* at the post-transcription level by binding to multiple binding sequences present in its CDS region, and promotes degradation by RNase III.

**Fig. 3:**
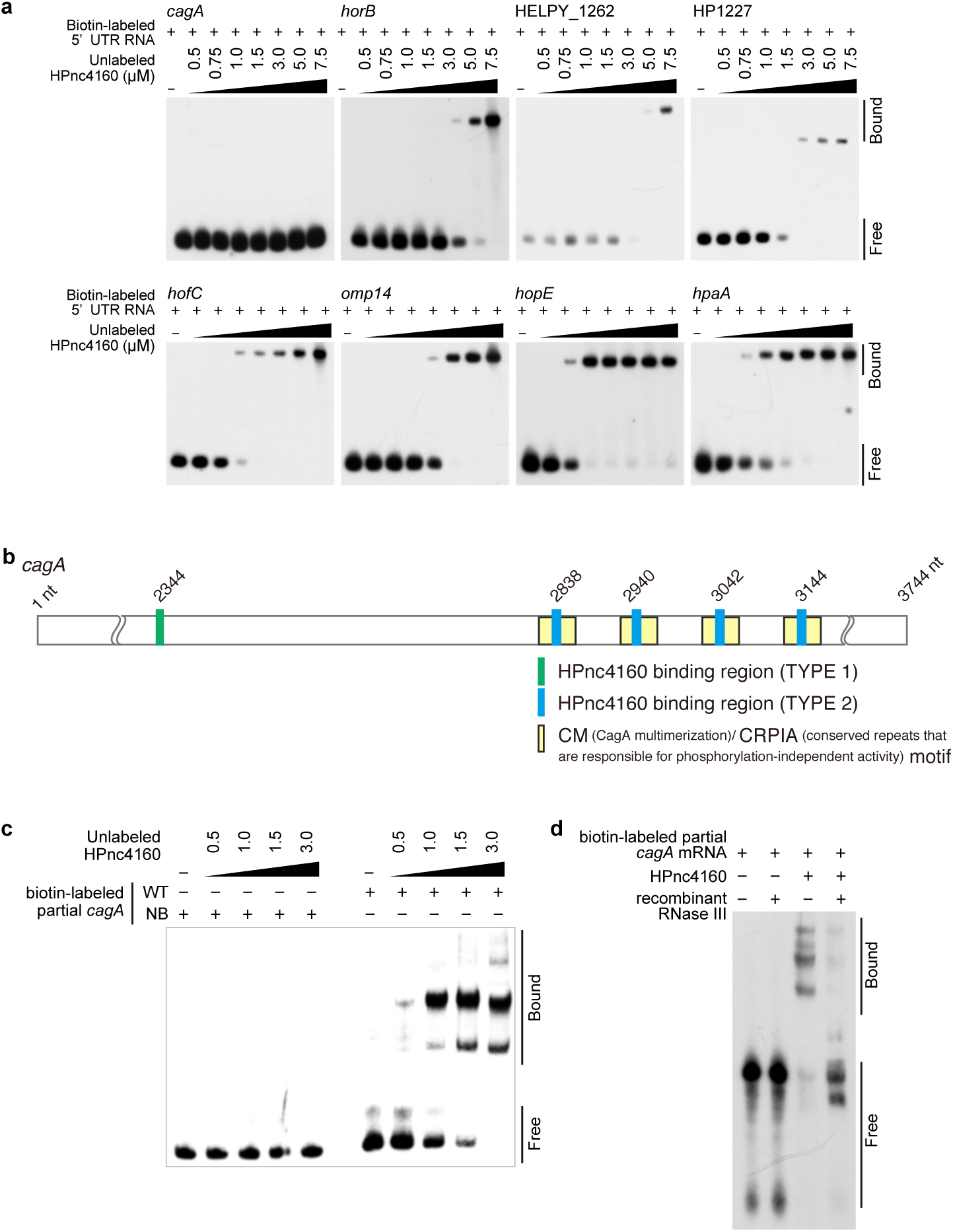
HPnc4160 binds to target mRNA. **a**, Electrophoretic mobility shift assay (EMSA) analysis of HPnc4160 binding to the 5’ UTR region of each candidate mRNA. **b**, Schematic of CagA motifs, and HPnc4160 binding sequences. **c**, EMSA analysis of HPnc4160 binding to RNA of partial *cagA* WT or HPnc4160-non-binding *cagA* (NB-*cagA*). **d**, RNase protection assay with HPnc4160, *cagA* mRNA, and recombinant RNase III.

## Effects of HPnc4160 on *H. pylori* pathogenicity

Among the factors that HPnc4160 regulated the expression levels, we further analyzed CagA, which has been deeply involved in pathogenesis. First, we confirmed whether the binding of HPnc4160-*cagA* mRNA actually controls the expression levels of *cagA* mRNA and protein in *H. pylori*. The quantitative PCR showed that, in the *H. pylor*i expressing NB-*cagA* in which all five HPnc4160-binding DNA sequences were mutated but the amino acid sequence was preserved (Extended Data Fig. 5g-h), the expression level of HPnc4160 was similar to that of the wild-type, but the expression level of *cagA* mRNA was significantly increased to the same extent as that of the Δ*hpnc4160*/*hpnc4170* strain (Extended Data Fig. 6a, Fig. 4a). Using the urease UreA protein as a loading control for *H. pylori*, we confirmed that NB-*cagA* strain expressed CagA protein at a higher level than wild-type and Δ*hpnc4160*/*hpnc4170* / pHel2-hpnc4160 strains, similar to Δ*hpnc4160*/*hpnc4170* strain (Fig. 4b). Next, we analyzed Western blot of the gastric epithelial cell line AGS infected with *H. pylori*. Using β-actin as a loading control for cells, we confirmed that the amount of UreA protein, which exhibited the bacterial amounts, was the same in the lysates of cells infected with any of the strains, indicating that all of the strains showed same binding ability to AGS cells (Extended Data Fig. 6b). Some of the CagA proteins injected from *H. pylori* to host epithelium via TFSS were tyrosine phosphorylated by host Src/Abl kinase and detected by pY-CagA-specific antibody. Using the antibody, we confirmed that the amount of intracellular CagA was increased in the NB-*cagA*-infected cells, accompanied with the increase in the amount of CagA (Extended Data Fig. 6b). Injected CagA induces AGS cell motility (scattering/hummingbird). In the AGS cells infected with the Δ*hpnc4160*/*hpnc4170* or the NB-*cagA* strains, more remarkably elongated cells were observed than in the wild-type or Δ*hpnc4160*/*hpnc4170* / pHel2-*hpnc4160* strain-infected cells (Fig. 4c-d). When we analyzed amount of IL-8 protein secreted from *H. pylori*-infected cells, which is induced mainly by intracellular CagA, we found that the *cagA*-NB strain infected cells had higher IL-8 producing ability than the wild-type infected cells (Extended Data Fig. 6c). These results suggested that binding of HPnc4160 to *cagA* mRNA is important for controlling the amount of functional CagA protein injected by *H. pylori*.

**Fig. 4:**
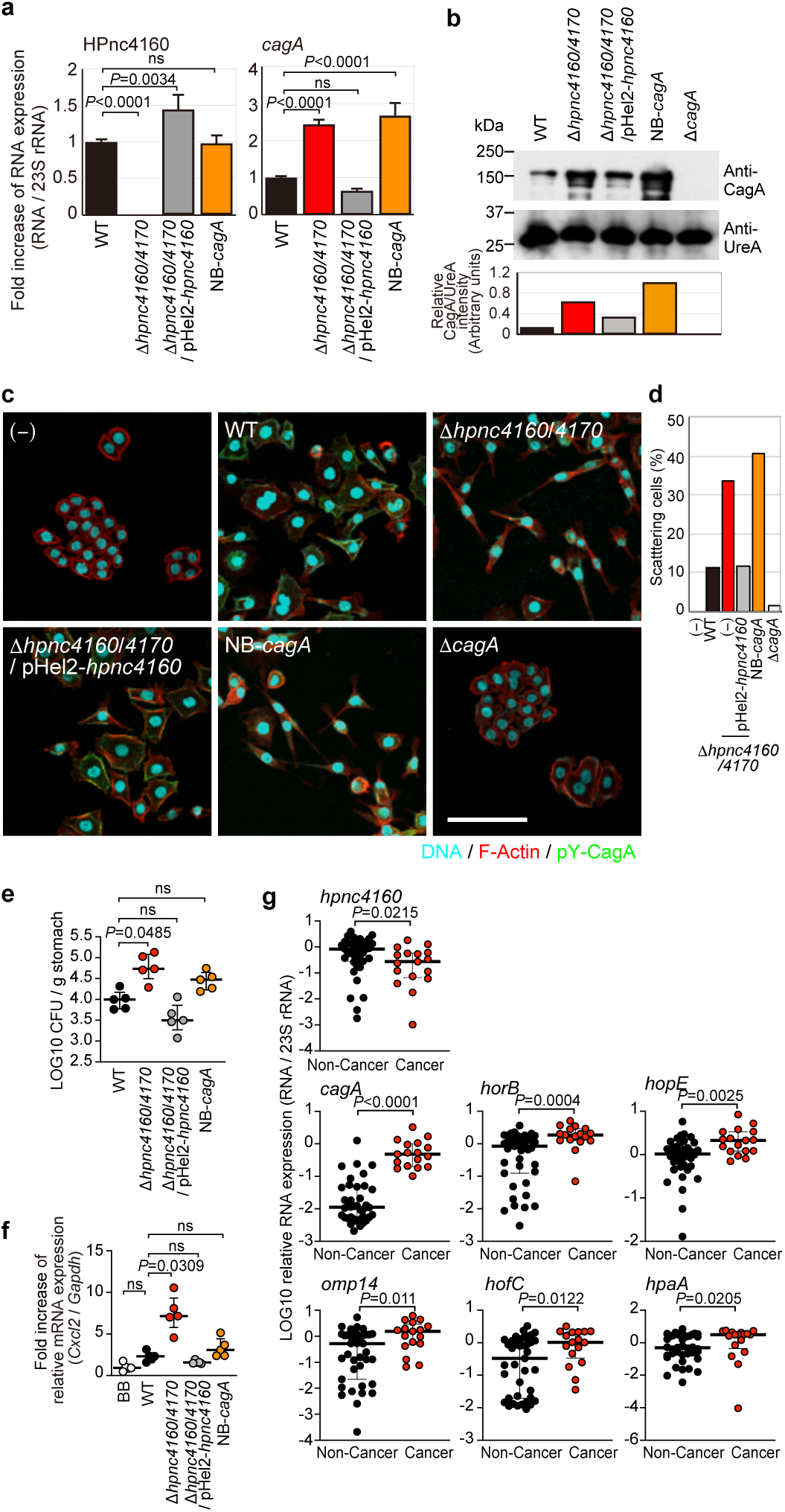
HPnc4160 controls bacterial host adaptation and pathogenesis. **a**, HPnc4160, and *cagA* mRNA expression levels. The results represent the average of three separate experiments (each n=3). Data are presented as means ± s.d. (error bars). *P* values are from non-parametric Dunnett’s multiple comparison test (at two-sided). ns: not significant. Experiments were repeated three times with similar results. **b**, Protein expression levels of CagA in each mutant strain. UreA protein levels serve as bacterial loading controls. **c** and **d**, Scattering phenotypes of *H. pylori*-infected AGS gastric epithelial cells. **c**, DNA (blue), F-actin (red), and anti-phosphorylated CagA antibody (pY-CagA, green) were stained. Scale bar, 50 μm. **d**, Quantification of scattering activity of AGS cells induced by *H. pylori* infection. **e** and **f**, HPnc4160 deletion mutants efficiently colonized the stomach and contributed to increasing mRNA levels of inflammatory chemokine *Cxcl2* in mice stomach. C57BL/6 mice were inoculated with *H. pylori*. At three days after infection, animals were sacrificed, and a quantitative culture assay (**e**) and a quantitative RT-PCR (**f**) were performed on gastric specimens. Data are median with interquartile range. *P* values are from non-parametric Dunn’s multiple comparison test (at two-sided). ns: not significant. **g,** Clinical isolates from malignant patients downregulate HPnc4160 and upregulate expression levels of its target genes. Expression levels of indicated mRNAs in clinical isolates of non-cancer (Non-Cancer, n=39) and cancer (Cancer, n=17) patients were quantified and normalized with the levels of 23S rRNA. Data are presented as medians with interquartile range. *P* values are from the non-parametric Mann-Whitney test (at two-sided).

To understand the significance of the HPnc4160 control mechanism in the bacterial adaptation to the host to establish infection, mice were orally inoculated with each strain, and the number of bacteria colonized in the stomach was analyzed three days post infection. The number of colonized bacteria in the stomach was significantly increased in the Δ*hpnc4160*/*hpnc4170* strain compared to the wild-type, but the Δ*hpnc4160*/*hpnc4170* / pHel2-*hpnc4160* and the NB-*cagA* strains were equivalent to wild-type (Fig. 4e). Since Δ*hpnc4160*/*hpnc4170* infection significantly increased *Cxcl2* mRNA compared to wild-type infection, but NB-*cagA* strain was equivalent to wild-type, it is suggested that factors other than CagA controlled by HPnc4160 may be important for the bacterial adaptation as well as development of gastritis (Fig. 4f). To confirm the significance of HPnc4160 in the pathogenesis of *H. pylori*, we examined the expression levels of the HPnc4160 target genes in clinical isolates. As shown in Fig. 4g, isolates from cancer patients had lower levels of *hpnc4160*, but increased expression of six factors controlled by HPnc4160 (*cagA*, *horB*, *hopE*, *omp14*, *hofC*, and *hpaA*), compared to isolates from non-stomach cancer patients (Fig. 4g, Extended data Fig. 7). These data strongly suggested that target mRNA expression was suppressed by HPnc4160 at the onset of *H. pylori* infection, and with the course of infection, thymidine repeats were inserted into the upstream region of HPnc4160, HPnc4160 expression decreased, and target mRNA expression increased; those episodes contributed to bacterial adaptation to the host environment, leading to gastritis and gastric cancer formation (Extended Data Fig. 8).

## Discussion

Among the bacterial factors, CagA had extremely high mRNA and protein levels in the bacterial cells (Fig. 2a and b). Our data indicated that T-stretch length in the upstream region of *hpnc4160* did not elongate under *in vitro* condition (Extended Data Fig. 3). However, the T-stretch length was increased and the expression levels of CagA and OMPs increased *in vivo* in gastric infection (Fig. 1e, 2a and 2b). Adaptation to the *in vitro* culture environment does not require CagA or OMPs. Therefore, under *in vitro* conditions, it may be that the expression level of hpnc4160 is increased and the expression of target is suppressed in order to allocate energy for the growth of bacterial cells, rather than expressing genes having high expression levels such as CagA. CagA suppresses apoptosis of the gastric mucosal epithelium and contributes to persistent infection of *H. pylori*^6^. When *H. pylor*i enter the stomach, the bacteria may have acquired a mechanism to decrease the expression of HPnc4160 to increase the expression of OMP and CagA simultaneously, in order to adapt to environmental changes to colonize the gastric mucosa for a scaffold for growth.

Gene expression control mechanism by variation of the number of repeat sequences is known as one of various gene expression control mechanisms by phase variation in *H. pylori*^21^. This study suggests that the repeat sequence of *H. pylori* genome is important not only as an ON / OFF mechanism of protein expression such as cell adhesion factors SabA and BabA, but also in sRNA expression. It has been reported that *H. pylori* DNA polymerase I does not have gene repairing activities like other bacteria, thus insertions or deletions of bases called slipped strand mispairing (SSM) occur in simple repetitive sequences^22^. In fact, since there were variations in poly-T length within the upward region of *hpnc4160* in *H. pylori in vivo* (Extended Data Fig. 2a), it is conceivable that *H. pylori* become genetically heterogeneous bacterial population using SSM during the course of infection, so that a population suitable for mutation in the host is selected and propagated. In this study we mainly used strain ATCC 43504, which is a clinical isolate originally from the human antrum. The poly-T length, which did not fluctuate in the *in vitro* subculture, increased in rodent isolates ranged from 14 to 19 copies (Extended Data Fig. 2, 3), indicating that the introduction of SSM may be caused by some host stress condition.

The HPnc4160-binding sequence in the ATCC 43504 *cagA* CDS was duplicated at five sites (Fig. 3b). In the regulation of mRNA expression by general sRNA, mRNA instability can be induced by binding at one site. This is the first report of a sRNA gene regulatory mechanism having multiple binding sequences in one gene of a pathogen. The four out of five HPnc4160-binding sequences in the *cagA* CDS was located in the CM/CRPIA motif of the *cagA* gene, which is involved in maintaining host epithelial cell structure^20^. Since the number of the CM/CRPIA motif differs depending on the strain, HPnc4160 may regulate differences in pathogenicity between Western and East Asian type of *H. pylori*.

The onset of the diseases due to *H. pylori* infection is thought to be the result of persistent infection for decades after the initial infection during childhood, when a very small number of bacteria-containing aerosol were taken into the body orally^23^. However, since it is difficult to analyze *H. pylori* infection experiments using laboratory animals for decades, a relatively short time infection analysis must be performed by inoculating a large number of bacteria. Therefore, the mutation analysis in this study may correspond to mutations acquired in the acute phase (Supplementary information 1 and 3)^15, 24^. Infection by a small number of bacteria that mimic natural infection allows for more detailed analysis of the establishment of infection.

In this study, we investigated the mutations of *H. pylori*, originated from the same strain, infected in experimental animals with the same genetic background and environment. Unlike studies in individuals with widely differing genetic backgrounds, stomach environments, and infection history, our study has advantage for understanding the adaptation process of *H. pylori* to the host. We discovered a novel non-coding sRNA that is important for post-transcriptional translation control of pathogenic factors of *H. pylori*, such as CagA, which was previously considered to be the most important pathogenic factor for gastric cancer development, and putative OMPs involved in bacterial adhesion. This study is not limited to elucidating the complicated mechanism of persistent *H. pylori* infection, but its application to *H. pylori*-specific therapies that do not rely on antibiotics can also be expected.

## Methods

### Data reporting

No statistical methods were used to predetermine sample size, and the experiments were not randomized and the investigators were not blinded to allocation during experiments and outcome assessment.

### Strains and culture conditions

The *Helicobacter pylori* strain ATCC 43504, its isogenic mutants Δ*cagA* and Δ*virB7*, strains SS1 and PMSS1 have been described previously^6, 25^. *H. pylori* was cultured on Trypticase soy agar with 5% (v/v) sheep blood (Thermo Fisher Scientific, Waltham, MA, USA) for 2 days at 37°C in microaerobic conditions. Bacterial colonies were suspended in Brucella broth (Thermo Fisher Scientific) supplemented with 5% (v/v) inactivated FBS (Thermo Fisher Scientific), adjusted to Optical density 600 nm of 0.05, and incubated 15 hours at 37°C with gentle agitation under microaerobic conditions.

The AGS human gastric epithelial cell line (ATCC CRL-1739) was maintained in DMEM/F-12 (Thermo Fisher Scientific) containing 10% (v/v) FBS. AGS cells were seeded in six-well plates and grown to ∼80% confluence to be used for western blot analysis. For immunofluorescence microscopy, cells were seeded in six-well plates with cover glass, and grown to ∼80% confluence.

### Antibodies and immunohistochemical reagents

The anti-Tyr(P)-CagA, and anti-UreA polyclonal antibodies have been described previously (Mimuro MC 2002). Anti-CagA polyclonal antibody was purchased from AUSTRAL Biologicals (CA, USA), anti-actin monoclonal antibody was from MERCK (Darmstadt, Germany), Horse radish peroxidase (HRP)-labeled anti-rabbit IgG and HRP-labeled anti-mouse IgG, and FITC-labeled anti-rabbit IgG was from Jackson ImmunoResearch Laboratories Inc. (PA, USA). DAPI was from SIGMA-ALDRICH (MD, USA), and Rhodamine Phalloidin was from Thermo Fisher SCIENTIFIC (MA, USA).

### Animal infection

*H. pylori* infection of rodents were performed as described previously^26^. Briefly, 6-week-old male MON/Jms/GbsSlc Mongolian gerbils were orally administered with 200 μL of Vancomycin (500 mg/L) at 24 and 48 hours before *H. pylori* inoculation. On the days of *H. pylori* inoculation, 300 μL of 5% (w/v) sodium bicarbonate were orally administrated 10 minutes before bacterial inoculation. The gerbils were then intragastrically inoculated with an *H. pylori* culture containing 10^9^ CFU for 2 consecutive days. As for C57BL/6 mice (SLC Japan Inc., Tokyo, Japan) were intragastrically inoculated once with *H. pylori* culture of 10^9^ CFU. After indicated date, the stomach of each infected animal was opened along the greater curvature. To quantitatively isolate *H. pylori,* the stomach was excised, weighed, and homogenized. Serial dilutions were plated on *H. pylori*-selective agar plates (Eiken Chemical Co.) and incubated under microaerophilic conditions at 37°C for 4 days, after which the cfu were counted. Colonization data points of 1 × 10^3^ cfu were the minimal detection limit of the assay.

For isolation of strains recovered from *H. pylori*-infected rodents, each colony on the *H. pylori*-selective agar plates were picked up and spread on Trypticase soy agar with 5% (v/v) sheep blood, and incubated under microaerophilic conditions at 37°C for two days. Then, the colonies were suspended in Brucella broth supplemented with 5% (v/v) inactivated FBS, adjusted to Optical density 600 nm of 0.05, and incubated 15 hours at 37°C with gentle agitation under microaerobic conditions. The cultures were preserved with 50% (v/v) glycerol in −80°C until use.

For RNA isolation, the tissue was immediately frozen in liquid nitrogen. Animal experiments were conducted in accordance with the University of Tokyo or Osaka University guidelines for the care and use of laboratory animals and were approved by the ethics committee for animal experiments at the University of Tokyo or Osaka University.

### Genomic DNA purification and sequencing

For PCR templates, genomic DNA was purified using InstaGene Matrix (Bio-Rad Laboratories, Inc., CA, USA).

For whole genome sequencing, genomic DNA was purified from mid-log phase culture of strain ATCC43504 using QIAGEN DNeasy (QIAGEN). A genomic DNA library for sequencing was prepared using the Nextera XT DNA Sample Preparation kit (Illumina, San Diego, CA, USA) and sequenced using the Illumina MiSeq (for isolates from gerbils) or HiSeq X (for isolates from mice) platform to generate 300-bp paired-end reads. Genome assembly, scaffolding, and gap-closing were performed using the Platanus assembler (Kajitani et al. 2014). Gene identification and annotation were conducted by the Microbial Genome Annotation Pipeline (MiGAP [http://www.migap.org]). The raw read sequences and assembled scaffold sequences have been submitted to the DDBJ/EMBL/Genbank under the Bioproject accession number; SAMD00178897-SAMD00178935, SAMD00179460, SAMD00178937 and SAMD00204457-SAMD00204466.

The DNA sequences mutated in more than 50% of the 40 strains recovered from Mongolian gerbils, or, in all of the 10 strains recovered from C57BL/6 mice were listed in Extended Data Table 1. We selected the genes to further analyze for their mRNA expression levels as follows. For the gene in which the mutation was in the CDS region, the mRNA expression level of the CDS was measured. While, when the mutation insertion region was an intergenic region, we measured the mRNA expression level of an adjacent gene in which the intergenic region could be a 5’UTR region. As for HP1243 and HPG27_298, which started from 3’ end of HP1243 with 33 nucleotides spaces, were regarded as a continuous gene; since both genes are annotated as *babA* gene and ribosomal binding site (RBS) is assigned only at the upstream region of HP1243.

### *In vitro* passage experiment

*H. pylori* ATCC 43504 was recovered from frozen stock and cultured on 5% (v/v) sheep blood agar for 2 days at 37°C in microaerobic conditions. Bacterial colonies were suspended in 3 tubes of Brucella broth supplemented with 5% (v/v) inactivated FBS. Each bacterial suspension was adjusted to Optical density 600 nm of 0.05, and incubated 12 hours at 37°C with gentle agitation under microaerobic conditions. Following this incubation, each fraction of the suspension was preserved by freezing in 50% (v/v) glycerol as “Original” strains. Meanwhile, each bacterial suspension was sub-cultured by resuspending in Brucella broth supplemented with 5% (v/v) inactivated FBS to adjust Optical density 600 nm of 0.05, and incubated additional 12 hours at 37°C with gentle agitation under microaerobic conditions. The sub-cultivation was repeated for 60 passages (30 days), and each cell suspension was preserved by freezing in 50% (v/v) glycerol as “60-passaged” strains. The “Original” and “60-passaged” strains were recovered from frozen stock on 5% (v/v) sheep blood agar by 2 days incubation under microaerobic conditions, and then the colonies were suspended in Brucella broth supplemented with 5% (v/v) inactivated FBS and incubated 12 hours at 37°C with gentle agitation under microaerobic conditions. The bacterial cells were collected and subjected to the genomic DNA purification.

### RT-PCR

For preparation of total RNA from *H. pylori*, the liquid cultures of *H. pylori* were agitated under microaerobic conditions at 37°C overnight until the OD value at 600 nm reached 0.9.

Total RNA was extracted using ISOGEN (Nippon Gene, Tokyo, Japan), according to the manufacturer’s instructions. The concentration of the purified total RNA was analyzed using the NanoDrop Spectrophotometer (ThermoFisher Scientific, Wilmington, DE, USA). The total RNA was reverse transcribed into cDNA with miScript II RT Kit (QIAGEN) according to the manufacturer’s instructions. The levels of mRNA expression were quantified and normalized to 23SrRNA (for *H. pylori*) or *Gapdh* (for mice) expression with a THUNDERBIRD SYBR qPCR (TOYOBO) using the primer pairs described in Supplementary Information 6. The results are expressed as the means ± SEM from triplicate strain experiments.

### Genetic manipulation

#### Construction of plasmids for producing gene-deficient mutants

Isogenic gene null mutants derived from ATCC 43504 were constructed by insertional mutagenesis as follows. Using the extracted *H. pylori* ATCC 43504 genome as a template, DNA fragments containing the upstream region 500 bp and the downstream region 500 bp of the target gene were amplified by PCR using primer (CagA KO up XhoI, CagA KO up EcoRI, CagA KO down BamHICagA KO down NotI, HPnc4160/4170 KO up KpnI, HPnc4160/4170 KO up ClaI, HPnc4160/4170 KO down BamHI, HPnc4160/4170 KO down SacI; listed in Supplementary Information 6). The DNA fragments were introduced at the both sides of the *aphA3* (which confers kanamycin resistant) in pBluescript II SK (+) plasmid. The fragments from the resulted plasmid were introduced into *H. pylori* by electroporation.

#### Construction of non-marker H. pylori mutants

For constructing non-marker *H. pylori* mutants, ATCC 43504 *flaA* and *cag1* promoter and terminator were cloned into pBluescript SK(+) *Sma*I aphA3 *Sma*I, and *sacB* gene was cloned into *EcoR*I site (pKSB plasmid).

Mid-log-phase (OD600 = 0.5-0.7) of *H. pylori* in 20 ml culture liquid were washed twice with ice-cold 10% glycerol and resuspended by 200 μl of ice-cold 10% glycerol. 1μg of pKSB vector containing aimed mutation and the bacterial liquid were mixed at 4℃ and electroporated by Micropulser (Bio-Rad) with Ec2 (2.5kV) setting. After 4 hours incubation at 37°C in microaerophilic condition, cells were plated on 5% sheep blood agar plate TSAII containing 4 μg/ml Kanamycin and incubated 2-3 days at 37°C under the microaerophilic condition. 4 single colonies were seeded on new 5% sheep blood agar plate TSAII supplemented with 4 μg/ml Kanamycin and incubated for additional 2 days. Each colony was picked up and were cultured in Brucella broth containing 5% FBS at 37°C under the microaerophilic condition until *H. pylori* were grown to mid-log phase. 100μl of the medium were plated on 5% sheep blood agar plate supplemented with 2.5% sucrose and cultured for 2 days. Each colony was seeded on a new 5% sheep blood agar plate without antibiotics and incubated for 2 days. At the same time, the colony was seeded on a different agar plate with 4 μg/ml Kanamycin to confirm the Kanamycin resistant was disappeared. Grown *H. pylori* were transferred to liquid culture and the genome sequence was confirmed by Sanger sequencing.

#### Construction of point mutated H. pylori

The *H. pylori* recombination plasmids to establish various mutant strains (T15mut, T16mut, T17mut, T18mut, T19mut) in the upstream region of *hpnc4160* were constructed by PCR using *H. pylori* genome DNA from the strains isolated from gerbil after 8 weeks as a template, and primers (pKSB-HPnc4160 Point mut *Apa*I and pKSB-HPnc4160 Point mut *Xho*I; listed in Supplementary Information 6), then, the resulted DNA fragments were cloned into suicide pKSB plasmid.

*H. pylori* T15mut, T16mut, T17mut, T18mut and T19mut mutants were established by introducing each pKSB-based plasmid into *H. pylori* ATCC 43504 strain.

#### Construction of NB-cagA-expressing H. pylori

Based on the full length cagA cDNA sequence of ATCC 43504, we designed HPnc4160-unbound *cagA* gene sequence (NB-*cagA*, Extended Data Fig. 5g and h). The NB-*cagA* cDNA were artificially synthesized as pEX-K4J2-cagA mutant of 908 bps (eurofins, 99900008281-1). The cDNA fragments containing mutated *cagA* sequence were amplified using primers (pKSB-CagA-NB-ApaI, pKSB-CagA-NB-XhoI, listed in Supplemented Information 6), and cloned into a suicide vector pKSB. The resulted plasmids were introduced to *H. pylori* ATCC 43504 to obtain NB-cagA-expressing *H. pylori*.

#### Construction of hpnc4160 over-expressing H. pylori

The plasmid for the hpnc4160 overexpressing strain in *H. pylori* was constructed by combination of DNA fragments of hpnc4160 regions were amplified by PCR using primers (pHel2-4160-de-4170-hed-f XhoI, pHel2-4160-de-4170-hed-r BamHI, Supplemented Information 6) and genome DNA of the ATCC 43504 strain as a template. The resulted DNA fragments included the upstream region of *hpnc4160* without including the 5’ region of the hpnc4170 region. The DNA was cloned into pHel2 shuttle vector, and introduced into *H. pylori* by electroporation.

### RNA-seq

*H. pylori* were agitation under aerobic conditions and cultured at 37°C overnight until the OD value at 600 nm reached 0.9. Total RNA from the *H. pylori* were extracted using RNeasy (QIAGEN), according to the manufacturer’s instructions. The concentration of total RNA extracted was examined using the NanoDrop Spectrophotometer (ThermoFisher Scientific, Wilmington, DE, USA), according to the manufacturer’s instructions. Ten micrograms from each total RNA sample were treated with the MICROBExpress Bacterial mRNA Enrichment kit (Ambion, Grand Island, NY, USA) and RiboMinus™ Transcriptome Isolation Kit (Bacteria) (Invitrogen, Grand Island, NY, USA) following the manufacturer’s instructions. Samples were resuspended in 15 μL of RNase-free water. Bacterial mRNAs were chemically fragmented to the size range of 200-250 bp using 1 × fragmentation solution (Ambion, Grand Island, NY, USA) for 2.5 min at 94°C. cDNA was generated according to instructions given in SuperScript Double-Stranded cDNA Synthesis Kit (Invitrogen, Grand Island, NY, USA). Briefly, each mRNA sample was mixed with 100 pmol of random hexamers, incubated at 65°C for 5 min, chilled on ice, mixed with 4 μL of First-Strand Reaction Buffer (Invitrogen, Grand Island, NY, USA), 2 μL of 0.1 M DTT, 1 μL of 10 mM RNase-freed NTPmix, 1 μL of SuperScript III reverse transcriptase (Invitrogen), and incubated at 50°C for 1 h. To generate the second strand, the following Invitrogen reagents were added: 51.5 μL of RNase-free water, 20 μL of second-strand reaction buffer, 2.5 μL of 10 mM RNase-free dNTP mix, 50 U E. coli DNA Polymerase, 5 U E. coli RNase H, and incubated at 16°C for 2.5 h. The Illumina Paired End Sample Prep kit was used for RNA-Seq library creation according to the manufacturer’s instructions as follows: Fragmented cDNA was end-repaired, ligated to Illumina adaptors, and amplified by 18 cycles of PCR. Paired-end 150-bp reads were generated by high-throughput sequencing with the Illumina Hiseq 2500 Genome Analyzer instrument. After removing the low-quality reads and adaptors, RNA-Seq reads were aligned to the corresponding ATCC 43504 genome using Tophat 2.1.0 (Trapnell et al 2009), allowing for a maximum of two mismatch. If reads mapped to more than one location, only the one showing the highest score was kept. Reads mapping to rRNA and tRNA regions were removed from further analysis. After getting the reads number from every sample, edgeR with TMM normalization method was used to determine the DEGs. Significantly differentially expressed genes (FDR value < 0.05 and at least two-fold changes) were selected for further analysis.

### iTRAQ

*H. pylori* ATCC 43504 strains of wild-type, Δ*hpnc4160*/*hpnc4170*, and Δ*hpnc4160*/*hpnc4170* / pHel2-*hpnc4160* were cultured in Brucella Broth containing 5% FCS to OD600 = 0.9. 1.5 mL of each bacterial solution was centrifuged at 5,000 xg for 10 minutes at 4°C. The pellet was resuspended in Wash buffer (1 M KCl, 15 mM Tris-HCl, pH 7.4), centrifuged again, and the supernatant was removed. The pellet was resuspended in a Wash buffer containing 1 mM AEBSF (4-(2-Aminoethyl) benzenesulfonyl fluoride hydrochloride) and frozen at −80°C. iTRAQ analysis was commissioned to Filgen Corporation.

### EMSA (electrophoretic mobility shift assay)

cDNA fragments of small RNA HPnc4160 whole region, the fragments of 150 bp total of each 5′UTR region [from 100 bases upstream from the ribosome binding region (RBS), to 50 bases downstream of the RBS] (hp0410 gene, hp0486 gene, *horB* gene, hp0671 gene, *hopE* gene, *cagA* gene, hp1227 gene and helpy_1262 gene), and cDNA of 459 bp total containing the hpnc4160-binding 4 region near the 3’ tail of the *cagA* gene, were amplified by PCR using primers (Small RNA HPnc4160 XhoI, Small RNA HPnc4160 EcoRI; HP0410 150bp XhoI, HP0410 150bp EcoRI; HELPY_0660 150bp XhoI, HELPY_0660 150bp EcoRI; HP0671 150bp XhoI, HP0671 150bp EcoRI; HP0486 150bp XhoI, HP0486 150bp EcoRI; HPSH_00635 150bp XhoI, HPSH_00635 150bp EcoRI; HPP12_0555 150bp XhoI, HPP12_0555 150bp EcoRI; HP1227 150bp XhoI, HP1227 150bp EcoRI; HELPY_1262 150bp XhoI, HELPY_1262 150bp EcoRI; CagA-B codding XhoI, CagA-B codding EcoRI; listed in Supplementary Information 6) and the ATCC43504 genome as a template. The PCR products were cloned into the position of the downstream of the T7 promoter region of the pBluescript SK (+) plasmid. The NB-*cagA* mutant RNA used in the gel shift assay was amplified with a T7 promoter by PCR using (T7 promoter CagA-NB EMSA PCR s, T7 promoter CagA-NB EMSA PCR as) as primers and synthesized pEX-K4J2-CagA mutant (eurofins, 99900008281-1) *cagA* as a template. The *cagA* mutant RNA were prepared in the same manner except for mutations in the HPnc4160-binding 4 region. RNA was transcribed from a DNA fragment using an *in vitro* Transcription T7 kit (TAKARA).

Gel shift assays were performed using 0.04 pmol of 3’-biotin-tagged mRNA with increasing amounts of purified small RNA HPnc4160 in 20 μL reactions. Briefly, RNA was denatured (10 min, 80°C) and cooled for 5 min on ice. Yeast tRNA 1μg (ThermoFisher SCIENTIFIC) was added to the labelled RNA and the reaction was filled up to 10 μL with Binding Buffer (10 mM HEPES pH 7.3, 1 mM MgCl_2_, 20 mM KCl, 5% glycerol). 10 μL of either labelled mRNA was added to the HPnc4160. The mixtures were incubated at room temperature for 20 min. Then the samples were mixed with 5 μL native loading buffer before loading on a pre-cooled native 6% poly-acryl amide (PAA), 0.5x TBE gel. Gels were run in 0.5x TBE buffer at 30 mA per gel for 2 hours^27^.

### Cleavage assays

The cDNA of 720 bps of *H. pylori rnase* III was amplified by PCR using primers (pGEX-6P-1 RNaseIII XhoI-f, pGEX-6P-1 RNaseIII NotI-r, listed in Supplemented Information 6) and template (genome DNA from ATCC 43504 strain). The cDNA was cloned into pGEX6P-1 vector (GE). *E. coli* BL21 transformed with the plasmids were subjected to shaking culture in LB broth containing 100 μg/ml ampicillin at 37°C with constant shaking at 200 rpm. Protein expression was induced with IPTG to a final concentration of 0.1 mM, at 4°C, for 4 hours. The bacteria were collected by centrifugation and the pellets were subjected to GST-fusion protein purification using Glutathione Sepharose 4B (GE) according to the manufacture’s instruction. The RNase III protein was excised by PreScission Protease according to the manufacturer’s instructions. The purified protein derived from 6.7 mL of the bacterial culture was developed by SDS-PAGE, and the gel was stained with CBB to confirm that no contaminants were observed in the final product. The protein concentration was determined by absorbance at 280nm.

Nuclease assays using RNase III was performed using purified *H. pylori* recombinant RNase III. The gel shift assay protocol described above was followed, except that RNase III-specific buffer (25mM Tris pH 7.5, 50 mM NaCl, 50 mM KCl, 10 mM MgCl_2_, 1 mM DTT) was used instead of Binding Buffer. 3’-biotin-tagged partial *cagA* mRNA was incubated on ice with either 5 μM of small RNA HPnc4160 for 20 min. RNase III was then added at a final concentration of 300 nM and the reactions were incubated for 1 min at 37°C. The samples were mixed with 5 μL of native loading buffer before loading on a pre-cooled native 6% PAA, 0.5x TBE gel^28^.

### ELISA

AGS cells were co-incubated with *H. pylori* at an MOI of 100 for 12, 24, 36 hours at 37°C in a 5% CO_2_ environment in 24 well plates. The supernatants were collected and stored at −30°C. Enzyme-linked immunosorbent assays (ELISAs) for human IL-8 were performed using the Human IL-8 ELISA Kit (ThermoFisher SCIENTIFIC) according to the manufacturer’s instructions. The results are expressed as the means ± SEM from triplicate experiments.

### Immunofluorescence microscopy

AGS cells were infected with *H. pylori* at an MOI of 100 for 6 hours at 37°C in a 5% CO_2_ environment. The cells were fixed with 4% (w/v) paraformaldehyde-PBS at room temperature for 10 min. The cells were then washed with TBS for 3 times, and blocked with Saponin buffer [10% (v/v) Blocking One (Nakalai, Japan) containing 0.2% (w/v) saponin] at 4°C for 60 min. Antibodies used for staining were DAPI, Rhodamine Phalloidin (Thermo Fisher SCIENTIFIC, MA, USA), pyCagA. Confocal laser scanning microscopy (CLSM) image acquisition was performed using a Zeiss LSM 800 confocal laser scanning microscope with ZEN 2.3 software (Carl Zeiss, Jena, Germany).

## Extended Data Figures and Tables Legends

**Extended Data Figure 1.**
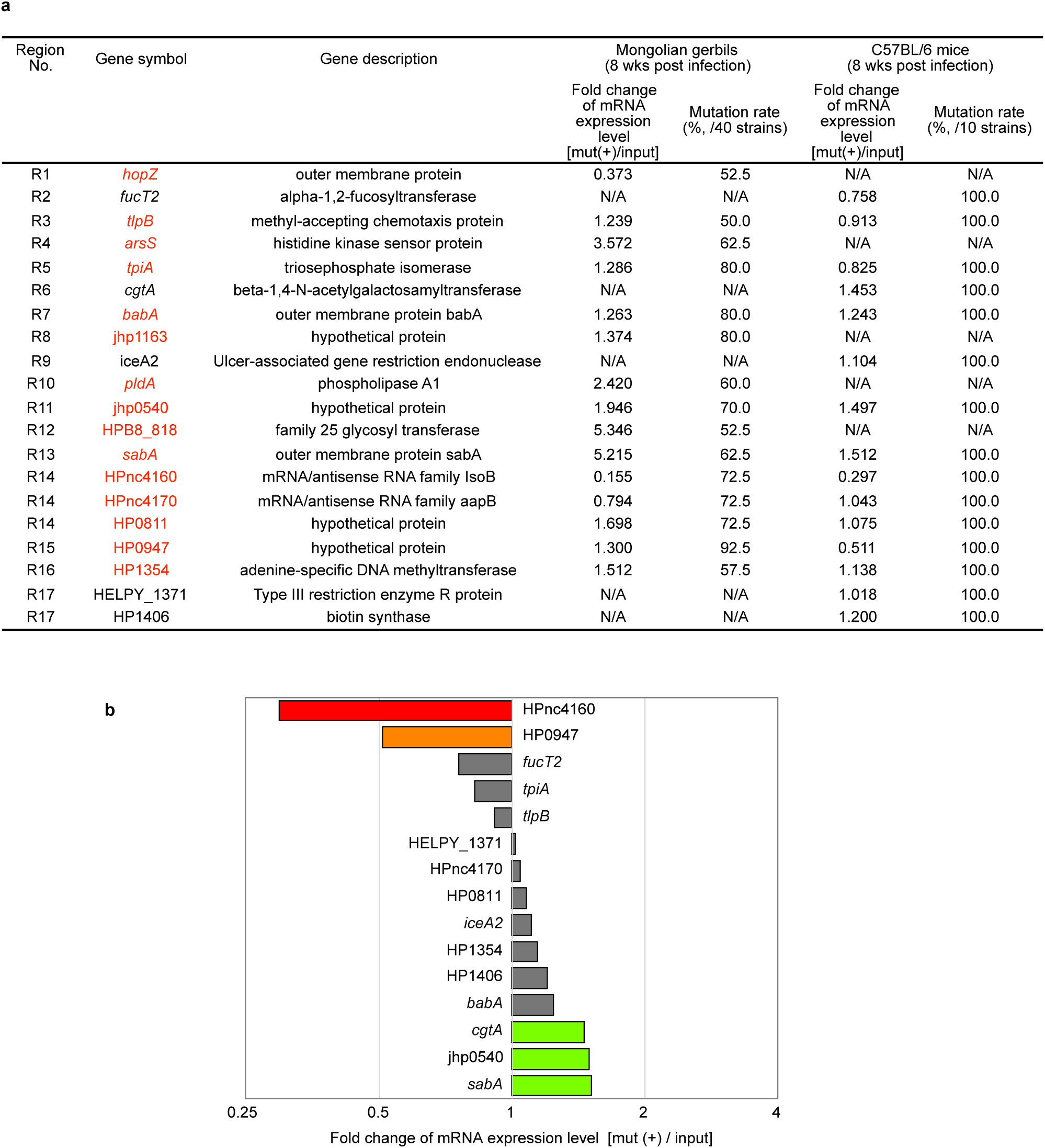
The expression levels of candidate RNAs of strains recovered from *H. pylori* ATCC43504-infected rodent stomachs. **a,** The list of mutation rates and expression levels of the candidate RNAs (mRNA or non-coding RNA) of strains recovered from stomachs of gerbils or mice 8 wks post-infection. The Locus tags highlighted in red indicated the candidates common in both of the strains originated from gerbils and mice. N/A, not applicable. **b,** The expression levels of candidate RNAs (mRNA or non-coding RNA) of isolates recovered from *H. pylori* ATCC43504-infected C57BL/6 mice stomachs. RNA expression levels of the genes or nearby genes of genome regions (Fig. 1b and Supplementary Information 2), which mutated in 100% of the recovered strains were assessed.

**Extended Data Figure 2.**
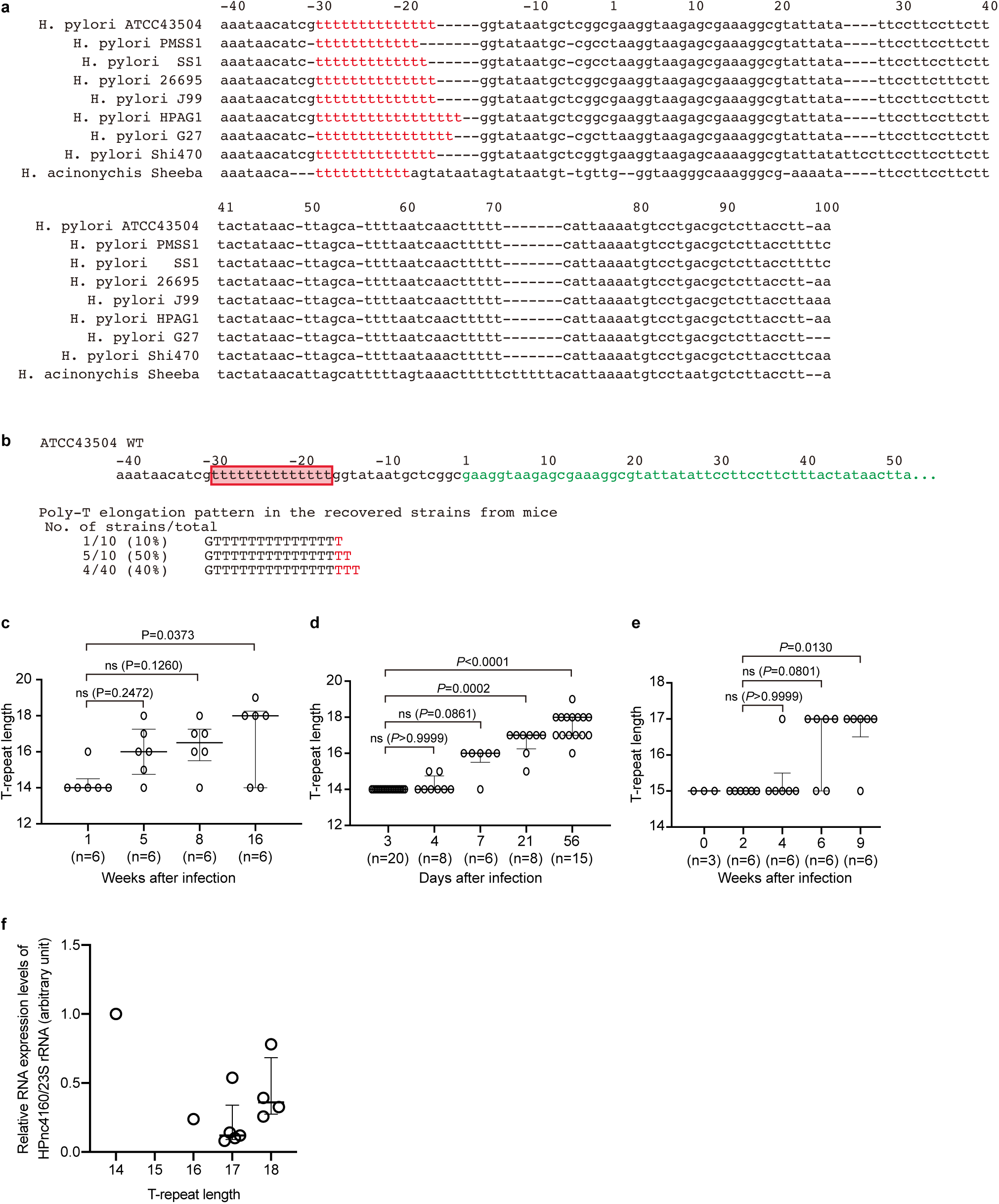
The length of the poly-T stretches upstream of the HPnc4160 coding region. **a,** Schematic diagram of genome DNA sequence around the HPnc4160 and poly-T sequence of genome analyzed strains. Red-colored “T” indicated stretches of poly-T sequence. **b,** Schematic diagram of genome DNA sequence around the HPnc4160 and poly-T sequence of mice-recovered strains 8 wks after infection (n=10, Hp1 to Hp10). Green indicated HPnc4160 transcribed sequence, red frame indicated poly-T mutated stretches in the upstream region of HPnc4160, and red-colored “T” indicated inserted nucleotides of each recovered strain compared with wild-type. **c-e,** Time-dependent change in the length of the poly-T stretches upstream of the HPnc4160 coding region in gerbils or mice-recovered strains. **c**, Strains from Mongolian gerbils infected with ATCC 43504. **d**, Strains from C57BL/6 mice infected with ATCC 43504. **e**, Strains from C57BL/6 mice infected with PMSS1. Data are median with interquartile range. *P* values are from non-parametric Dunn’s multiple comparison test (at two-sided). ns: not significant. **f**, Expression levels of HPnc4160 in the recovered *H. pylori* strains from mice (n=10) and *H. pylori* wild-type (T-repeat 14). Relative expression levels of HPnc4160 were measured by real-time PCR and plotted against T-repeat length in the upstream region of HPnc4160. Data are presented as means with s.d. *P* values represent the results of two-tailed Dunn’s multiple comparison test.

**Extended Data Figure 3.**
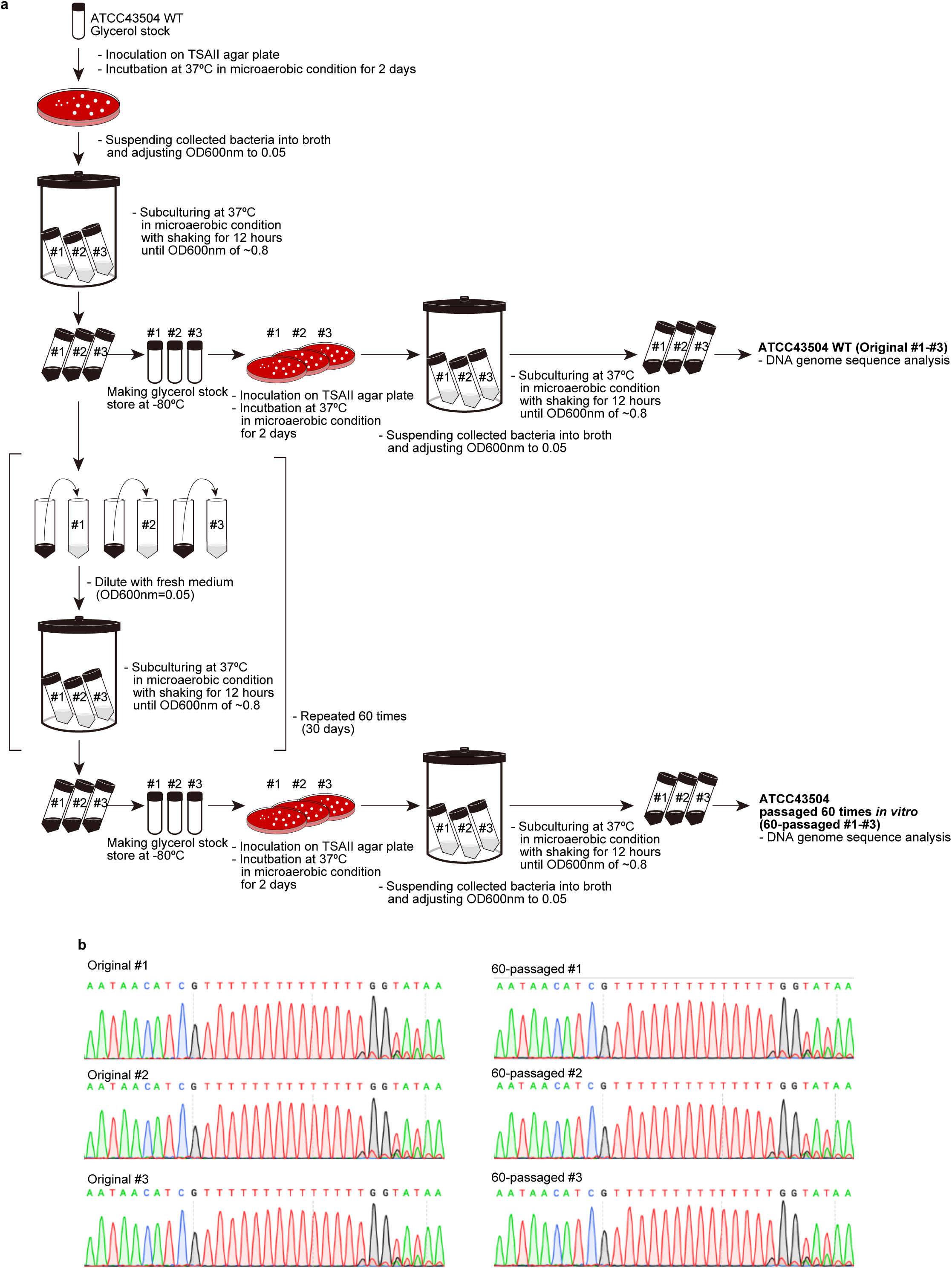
Effect of *in vitro* cultivation on the length of the poly-T stretches upstream of the HPnc4160 coding region. **a**, Experimental strategies schematic. **b**, The raw data of the DNA sequence analysis of *H. pylori* genomes prepared from original culture (Original #1 - #3) and from passaged *in vitro* for 60 times (60-passaged #1 - #3).

**Extended Data Figure 4.**
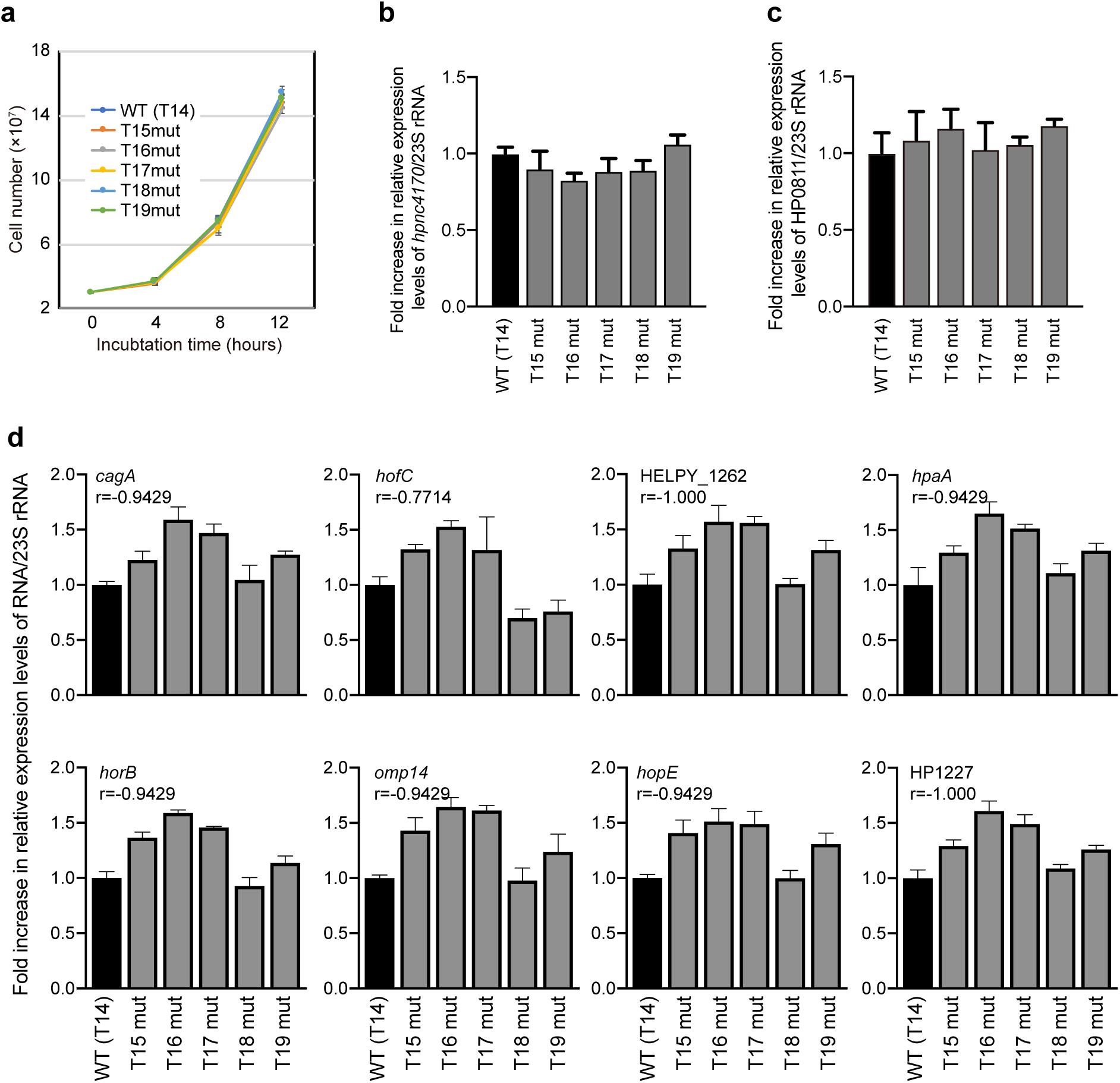
The length of the poly-T stretches upstream of the HPnc4160 coding region and RNA expression levels. **a**, Growth curves of *H. pylori* ATCC43504 mutants mutated in the number of T repeat in HPnc4160 upstream region. **b-c,** The relative expression levels of *hpnc4170* (**b**) and HP0811 (**c**) in the *H. pylori* strains genetically modified with the T-repeat length. Data are presented as means with s.d. (n=3). *P* values are from Dunnett’s multiple comparison test (at two-sided). ns: not significant. **d**, The relative RNA expression levels of target candidates of HPnc4160 showed an inverse correlation with HPnc4160. The total RNA from the indicated *H. pylori* ATCC 43504 strains were extracted, reverse transcribed, and provided for qPCR to assess the indicated genes. The results represent the average of three separate experiments (each n=3). Data are presented as means ± s.d. (error bars). Spearman correlation coefficients (r) were used to evaluate the relationships among relative RNA expression of HPnc4160 (Fig. 1h) and each target.

**Extended Data Figure 5.**
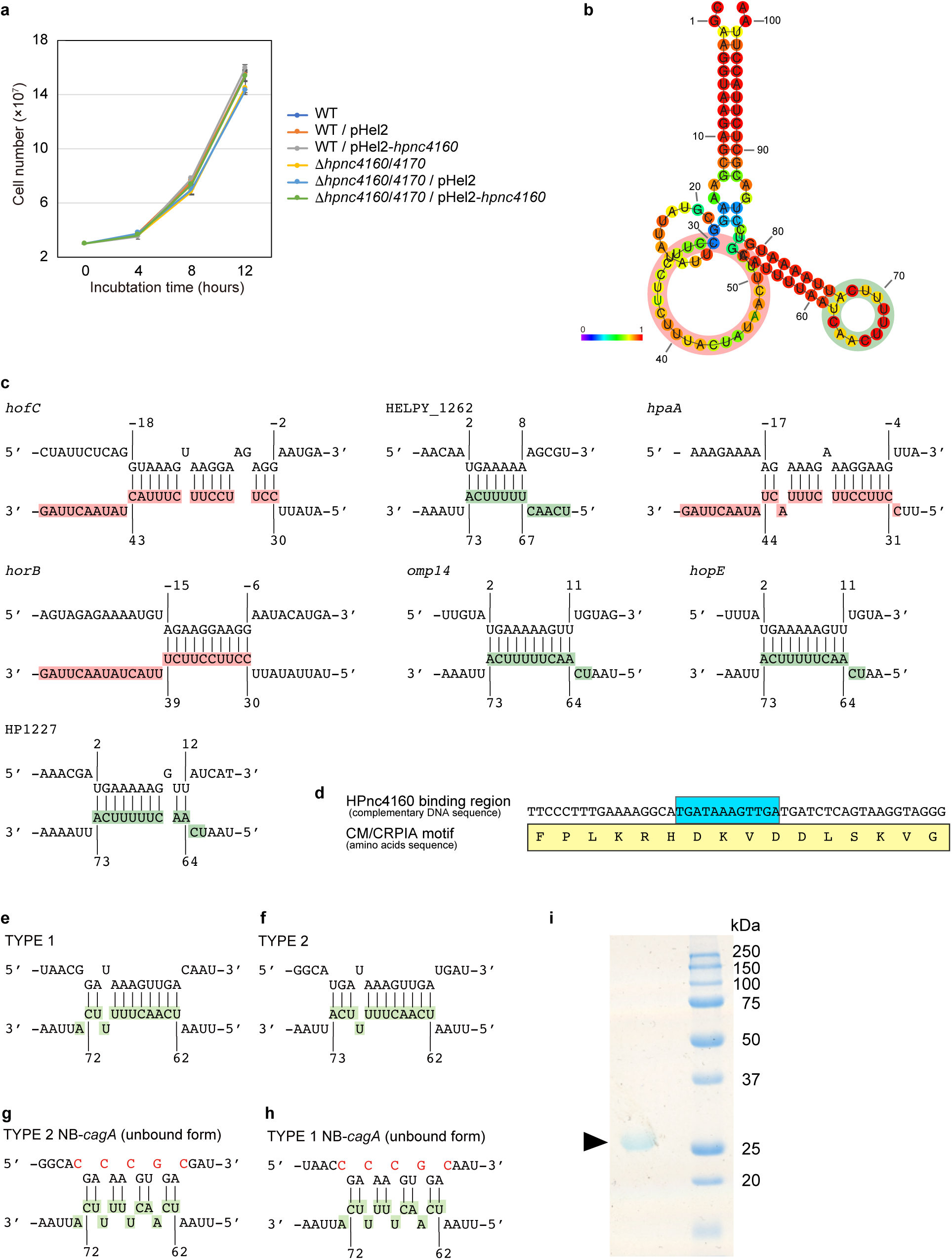
Predicted HPnc4160 binding sites. **a,** Growth curves of *H. pylori* ATCC43504 mutants. **b,** Predicted the secondary structure of HPnc4160 RNA by CentroidFold. The bases in the predicted structure are colored according to base-pairing probabilities. Circles in pink and light green color indicated loop structures having probabilities of binding to target RNA sequences. **c,** Schematic diagram of predicted HPnc4160 binding sites in the corresponding 5’UTR sequence of target genes. Upper sequences indicate target mRNA sequences with base numbers, whereas lower sequences indicate HPnc4160 sequence. Colored sequences are corresponding to the loop structures indicated in (**b**). **d**, Binding prediction of HPnc4160 and 5’ UTR of *cagA* mRNA. **d**, Schematic diagram of predicted HPnc4160 binding sites in the *cagA* CM/CRPIA motif of cagA CDS. e-f, Schematic diagram of predicted HPnc4160 binding sites in the corresponding CDS sequence of *cagA* TYPE 1 (**e**), TYPE 2 (**f**), and *cagA* nonbinding form (NB-*cagA*) of TYPE 2 (**g**) and TYPE 1 (**h**). Upper sequences indicate target *cagA* mRNA sequences, whereas lower sequences indicate HPnc4160 sequence with base numbers. Colored sequences are corresponding to the loop structures indicated in (**b**). Mutated nucleotides in *cagA* mRNA sequence are shown in red. **i**, Purified RNase III was separated by SDS-PAGE and stained with CBB.

**Extended Data Figure 6.**
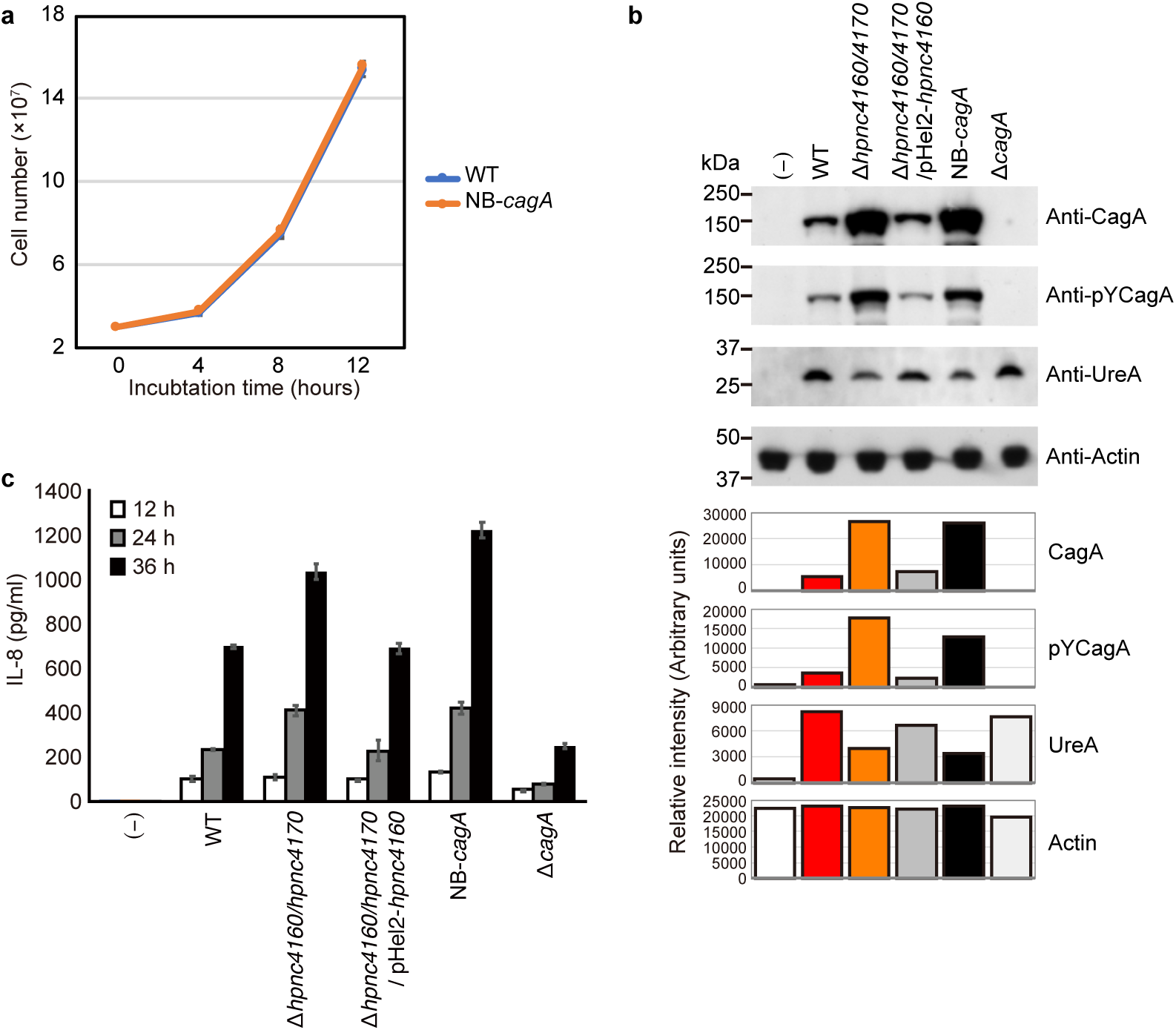
Effect of *cagA*-NB on host-cell-translocated CagA activity. **a,** Growth curves of *H. pylori* ATCC43504 *cagA*-NB mutant compared with wild-type. **b**, Phosphorylated CagA protein levels in cell lysates of AGS cells infected with *H. pylori* ATCC43504. The whole-cell lysates of AGS cells infected with *H. pylori* strains for 6 hours were subjected to western blot against anti-CagA, anti-pY CagA, anti-UreA, and anti-Actin antibodies. The band intensities were measured and calculated by ImageJ software. **c**, IL-8 production from AGS cells infected with *H. pylori* ATCC43504. The supernatants from AGS cells infected with *H. pylori* strains shown in the figure for the indicated time were subjected to ELISA system for IL-8 production. The results represent the average of three separate experiments (each n=3). Data are presented as means ± s.d. (error bars).

**Extended Data Figure 7.**
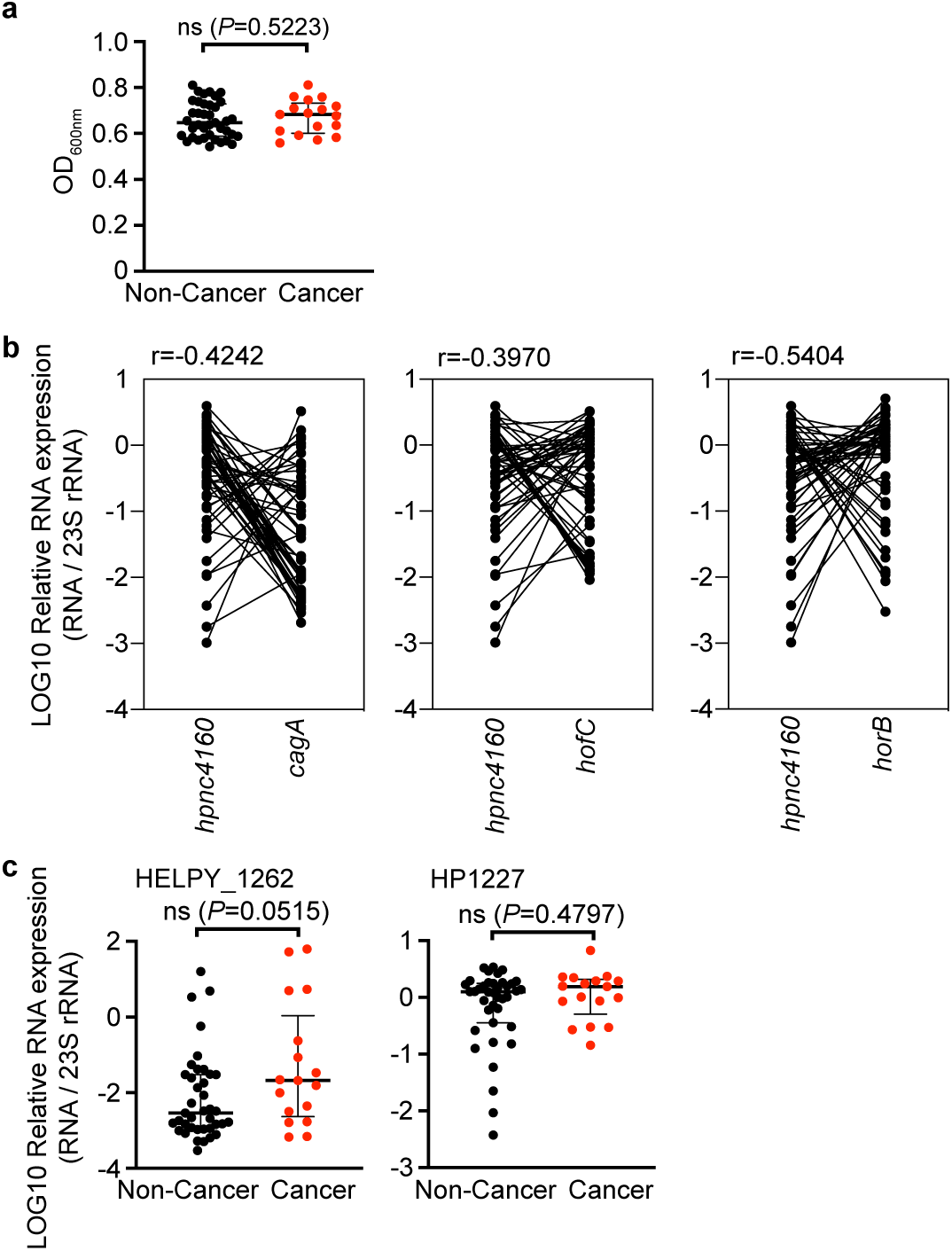
Characterization of clinical isolates. **a,** Clinical isolates of non-cancer (Non-Cancer, n=39) and cancer (Cancer, n=17) patients, which used in Fig. 4g, showed equal growth rate. The strains cultured on TSAII containing 5% sheep blood plates for 2 days were inoculated in Brucella broth containing 5% FCS, adjusted OD_600nm_ at 0.1, then cultured in microaerobic condition with agitation for 16 hours. The turbidity of the cultures was assessed at OD_600nm_. The Data are presented as medians with interquartile range. *P* values represent the results of the two-tailed Mann-Whitney test. ns: not significant. **b**, The relative RNA expression levels of target candidates of HPnc4160. Spearman correlation coefficients (r) were used to evaluate the relationships among relative RNA expression of HPnc4160 and each target. **c**, Comparison of expression levels of mRNA (HELPY_1262 and HP1227) in clinical isolates of non-cancer (NC, n=39) and cancer (C, n=17) patients. The expression levels of mRNA were normalized with the levels of 23S rRNA. Data are presented as medians with interquartile range. *P* values represent the results of the two-tailed Mann-Whitney test. ns: not significant.

**Extended Data Figure 8.**
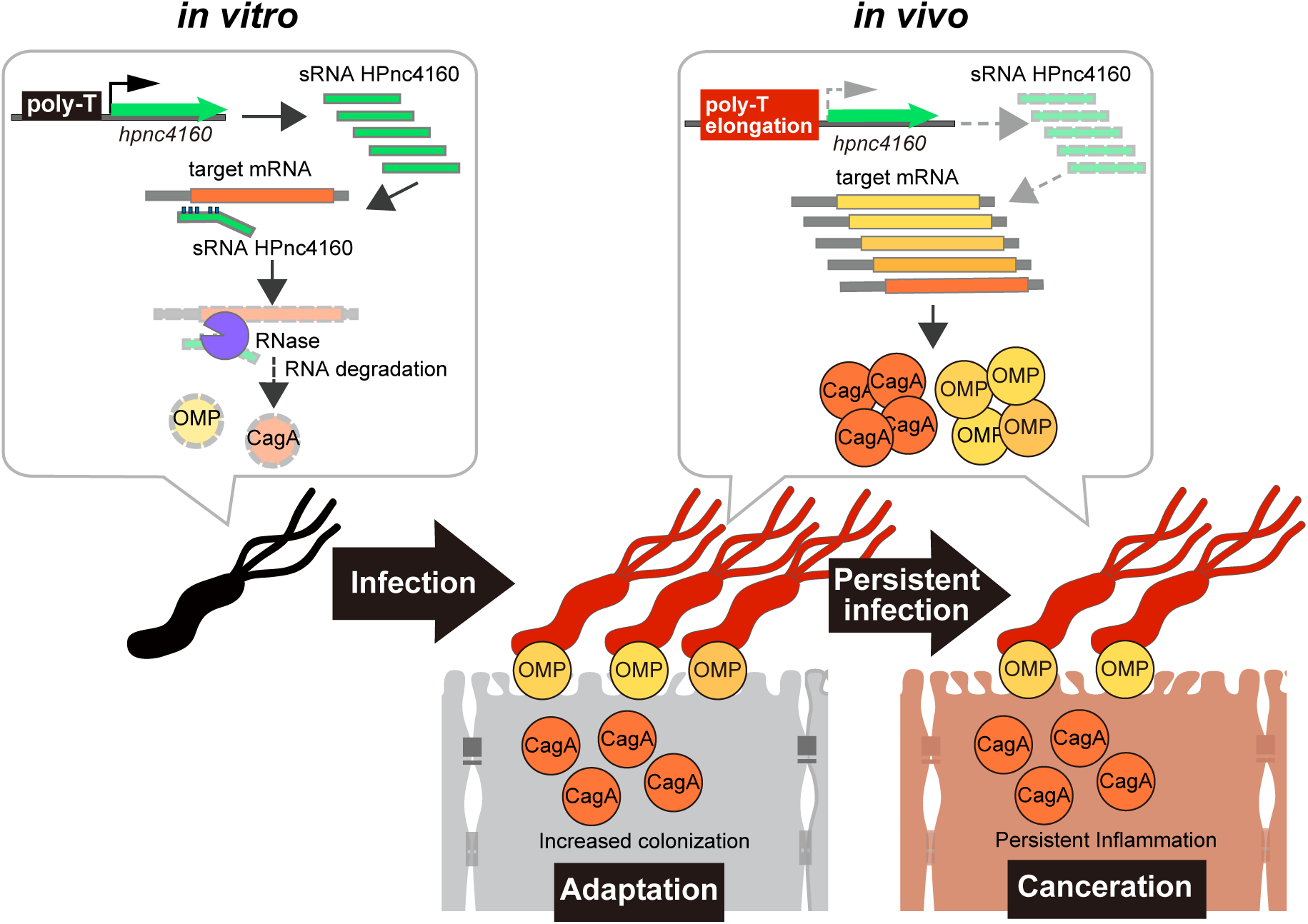
Infection-induced silencing of HPnc4160 upregulates target genes expression and promote bacterial host adaptation and canceration during chronic *H. pylori* infection. *H. pylori* infection *in vivo* leads elongation of T-stretch in the upstream region of HPnc4160 sRNA coding region, which results in decreased expression levels of sRNA HPnc4160. Gene silencing of HPnc4160 results in increased levels of target genes coding OMPs and CagA, and as a result, the levels of bacterial colonization and CagA translocation into the attached host cells were increased.

**Extended Data Table 1.**
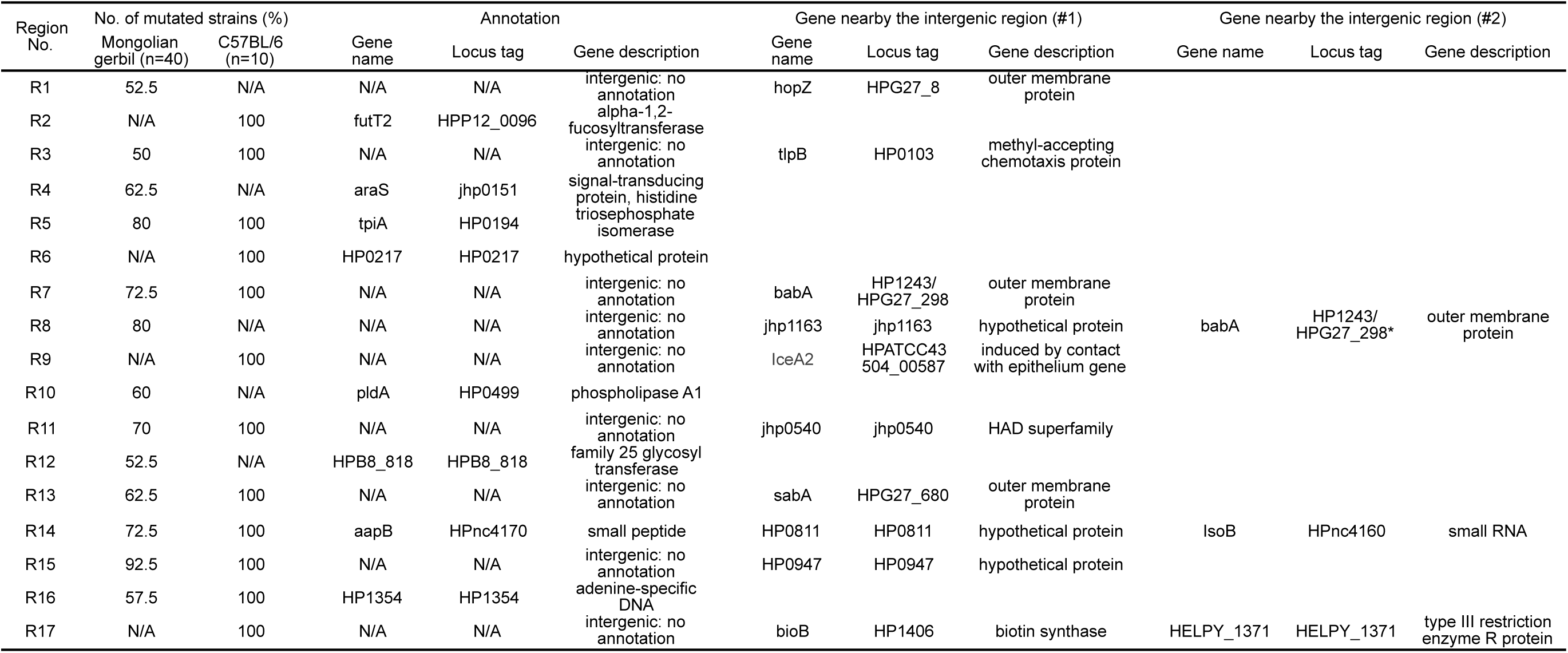
The list of mutated genome regions in the strains recovered from *H. pylori*-infected rodents’ stomachs. The list showed the genome regions that mutated in the strains isolated from the stomachs of rodents 8 weeks post infection. The DNA sequences in the regions listed in the table were mutated in more than 50% of the 40 strains recovered from Mongolian gerbils, or, in all of the 10 strains recovered from C57BL/6 mice. N/A, not applicable.

**Extended Data Table 2.** Comparative analysis of expression levels between *H. pylori* ATCC43504 Δ*hpnc4160*/*hpnc4170* mutant and wild-type strains. **a**, Comparative analysis of RNA expression levels between *H. pylori* ATCC 43504 Δ*hpnc4160*/*hpnc4170* mutant and wild-type strains by RNA-seq. Footnote | Normalized expression level and fold change of the strains were listed. Genes with *P*-values by Empirical Analysis of Digital Gene Expression in R (edgeR) test showed less than 0.001 were listed (17 factors). Eight genes selected by RNA-seq and iTRAQ analysis (Fig. 2c) were highlighted in red. **b**, Comparative analysis of protein expression levels between *H. pylori* ATCC43504 Δ*hpnc4160*/*hpnc4170* mutant and wild-type strains by iTRAQ. Footnote | Proteins showing relative protein abundance with *P*-value of less than 0.01 were listed (21 factors). Eight proteins selected by RNA-seq and iTRAQ analysis (Fig. 2c) were highlighted in red.

**a**
| Gene name | Average LOG2<br>normalized expression | LOG2 fold change<br>($\Delta hpnc4160/4170$ / WT) |
| --- | --- | --- |
| *cagA* | 15.6679564 | 0.8345757 |
| *vacA3* | 13.0181033 | 1.3000122 |
| *ureI* | 12.8122441 | 0.7256751 |
| *hofC* | 12.7562529 | 1.2385235 |
| *HP1227* | 11.1439545 | 0.5356426 |
| *hpaA* | 10.9856930 | 1.3994257 |
| *horB* | 10.9589406 | 1.0351050 |
| *omp14* | 10.8268799 | 1.3977712 |
| *hopE* | 10.8143652 | 1.0951572 |
| *ureE* | 10.6269817 | -1.1164069 |
| *HELPY_1262* | 10.6234286 | 0.9346910 |
| pAL226p12 | 9.6669233 | 0.8237894 |
| HP0487 | 9.1803234 | 1.7213025 |
| HELPY_0813 | 8.5854796 | 0.8765494 |
| *fliK* | 7.2801813 | -0.3748484 |
| *mraW* | 7.0680550 | 2.0913744 |
| HPAG1_1315 | 7.0164749 | -1.5753123 |

**b**
| Gene name | Fold change (LOG2,<br>$\Delta hpnc4160/hpnc4170$ / WT) | Pval (LOG10) |
| --- | --- | --- |
| *CagA* | 0.6831125 | -16.0000000 |
| *HofC* | 1.1079556 | -14.8875441 |
| *HpaA* | 1.4153801 | -14.5034266 |
| GroL | 0.2561653 | -10.0187654 |
| *HELPY_1262* | 0.6268596 | -9.9953361 |
| *HopE* | 1.1672937 | -7.7135456 |
| HP1409 | 0.1735112 | -3.6271647 |
| CeuE | 0.1737671 | -3.2477470 |
| NrdA | -0.1246917 | -2.8700087 |
| *HorB* | 0.8583789 | -2.6608050 |
| DnaK | 0.1192898 | -2.6579389 |
| GroES | 0.3291810 | -2.6528395 |
| Rpl9 | 0.2299570 | -2.6372507 |
| HP0305 | 0.2674759 | -2.5150013 |
| HopQ | 0.1519887 | -2.4589011 |
| *Omp14* | 1.1557492 | -2.4194920 |
| FlaB | -0.1694201 | -2.4003455 |
| Tuf | 0.1548424 | -2.3314305 |
| RpoB | -0.0584429 | -2.2822269 |
| Lpp20 | 0.1944653 | -2.1385174 |
| *HP1227* | 0.3642362 | -2.0521036 |

## Supplementary Information Legends

**Supplementary information 1.** Summary of mutations in the isolates recovered from *H. pylori*-infected Mongolian gerbils. The number of mutations in the isolates of 40 strains recovered from *H. pylori-*infected Mongolian gerbils’ stomachs 8 weeks after post-infection were listed.

| Animal No. | Strain Name | Total No. of mutations | SNPs (single nucleotide polymorphisms) |  |  |  | indel (insertion/deletion) |  |  | SNPs rate per base per year | indel rate per base per year |
| --- | --- | --- | --- | --- | --- | --- | --- | --- | --- | --- | --- |
|  |  |  | total | nonsynonymous | synonymous | intergenic | Total | genic | intergenic |  |  |
| #1 | S41 | 50 | 37 | 15 | 18 | 4 | 13 | 5 | 8 | 1.43E-04 | 5.04E-05 |
|  | S42 | 48 | 40 | 15 | 20 | 5 | 8 | 2 | 6 | 1.55E-04 | 3.10E-05 |
|  | S43 | 44 | 38 | 12 | 21 | 5 | 6 | 4 | 2 | 1.47E-04 | 2.33E-05 |
|  | S44 | 46 | 41 | 16 | 21 | 4 | 5 | 1 | 4 | 1.59E-04 | 1.94E-05 |
| #2 | S45 | 48 | 39 | 15 | 17 | 7 | 9 | 2 | 7 | 1.51E-04 | 3.49E-05 |
|  | S46 | 47 | 38 | 22 | 14 | 2 | 9 | 1 | 8 | 1.47E-04 | 3.49E-05 |
|  | S47 | 46 | 36 | 15 | 15 | 6 | 10 | 2 | 8 | 1.40E-04 | 3.88E-05 |
|  | S48 | 48 | 41 | 20 | 16 | 5 | 7 | 2 | 5 | 1.59E-04 | 2.71E-05 |
| #3 | S49 | 17 | 12 | 7 | 3 | 2 | 5 | 4 | 1 | 4.65E-05 | 1.94E-05 |
|  | S50 | 26 | 18 | 8 | 8 | 2 | 7 | 3 | 4 | 6.98E-05 | 2.71E-05 |
|  | S51 | 17 | 11 | 7 | 2 | 2 | 6 | 3 | 3 | 4.27E-05 | 2.33E-05 |
|  | S52 | 20 | 13 | 7 | 2 | 4 | 7 | 4 | 3 | 5.04E-05 | 2.71E-05 |
| #4 | S53 | 27 | 16 | 7 | 6 | 3 | 11 | 5 | 6 | 6.20E-05 | 4.27E-05 |
|  | S54 | 15 | 10 | 4 | 5 | 1 | 5 | 1 | 4 | 3.88E-05 | 1.94E-05 |
|  | S55 | 24 | 14 | 5 | 5 | 4 | 10 | 3 | 7 | 5.43E-05 | 3.88E-05 |
|  | S56 | 25 | 13 | 5 | 6 | 2 | 12 | 4 | 8 | 5.04E-05 | 4.65E-05 |
| #5 | S57 | 25 | 15 | 5 | 7 | 3 | 10 | 4 | 6 | 5.82E-05 | 3.88E-05 |
|  | S58 | 29 | 15 | 5 | 5 | 5 | 14 | 5 | 9 | 5.82E-05 | 5.43E-05 |
|  | S59 | 23 | 13 | 5 | 6 | 2 | 10 | 2 | 8 | 5.04E-05 | 3.88E-05 |
|  | S60 | 27 | 16 | 5 | 7 | 4 | 11 | 3 | 8 | 6.20E-05 | 4.27E-05 |
| #6 | S61 | 29 | 20 | 8 | 8 | 4 | 9 | 2 | 7 | 7.76E-05 | 3.49E-05 |
|  | S62 | 35 | 25 | 10 | 12 | 3 | 10 | 2 | 8 | 9.69E-05 | 3.88E-05 |
|  | S63 | 25 | 16 | 6 | 5 | 5 | 9 | 3 | 6 | 6.20E-05 | 3.49E-05 |
|  | S64 | 36 | 28 | 14 | 10 | 4 | 8 | 2 | 6 | 1.09E-04 | 3.10E-05 |
| #7 | S65 | 50 | 39 | 14 | 21 | 4 | 11 | 5 | 6 | 1.51E-04 | 4.27E-05 |
|  | S66 | 47 | 39 | 13 | 22 | 4 | 8 | 5 | 3 | 1.51E-04 | 3.10E-05 |
|  | S67 | 55 | 44 | 15 | 22 | 7 | 11 | 4 | 7 | 1.71E-04 | 4.27E-05 |
|  | S68 | 54 | 44 | 14 | 23 | 7 | 10 | 4 | 6 | 1.71E-04 | 3.88E-05 |
| #8 | S69 | 52 | 41 | 21 | 17 | 3 | 11 | 3 | 8 | 1.59E-04 | 4.27E-05 |
|  | S70 | 61 | 52 | 17 | 30 | 5 | 9 | 2 | 7 | 2.02E-04 | 3.49E-05 |
|  | S71 | 41 | 33 | 15 | 14 | 4 | 7 | 1 | 6 | 1.28E-04 | 2.71E-05 |
|  | S72 | 39 | 34 | 15 | 16 | 3 | 5 | 0 | 5 | 1.32E-04 | 1.94E-05 |
| #9 | S73 | 43 | 37 | 17 | 15 | 4 | 6 | 2 | 4 | 1.43E-04 | 2.33E-05 |
|  | S74 | 79 | 70 | 32 | 31 | 7 | 9 | 2 | 7 | 2.71E-04 | 3.49E-05 |
|  | S75 | 71 | 63 | 29 | 26 | 8 | 8 | 4 | 4 | 2.44E-04 | 3.10E-05 |
|  | S76 | 43 | 34 | 15 | 17 | 2 | 9 | 3 | 6 | 1.32E-04 | 3.49E-05 |
| #10 | S77 | 45 | 36 | 8 | 6 | 21 | 9 | 3 | 6 | 1.40E-04 | 3.49E-05 |
|  | S78 | 41 | 34 | 9 | 5 | 20 | 7 | 2 | 5 | 1.32E-04 | 2.71E-05 |
|  | S79 | 41 | 32 | 5 | 6 | 21 | 9 | 3 | 6 | 1.24E-04 | 3.49E-05 |
|  | S80 | 41 | 32 | 7 | 5 | 20 | 9 | 3 | 6 | 1.24E-04 | 3.49E-05 |
| Average |  |  |  |  |  |  |  |  |  | 1.19E-04 | 3.38E-05 |
| SD |  |  |  |  |  |  |  |  |  | 5.60E-05 | 8.51E-06 |

**Supplementary information 2.**
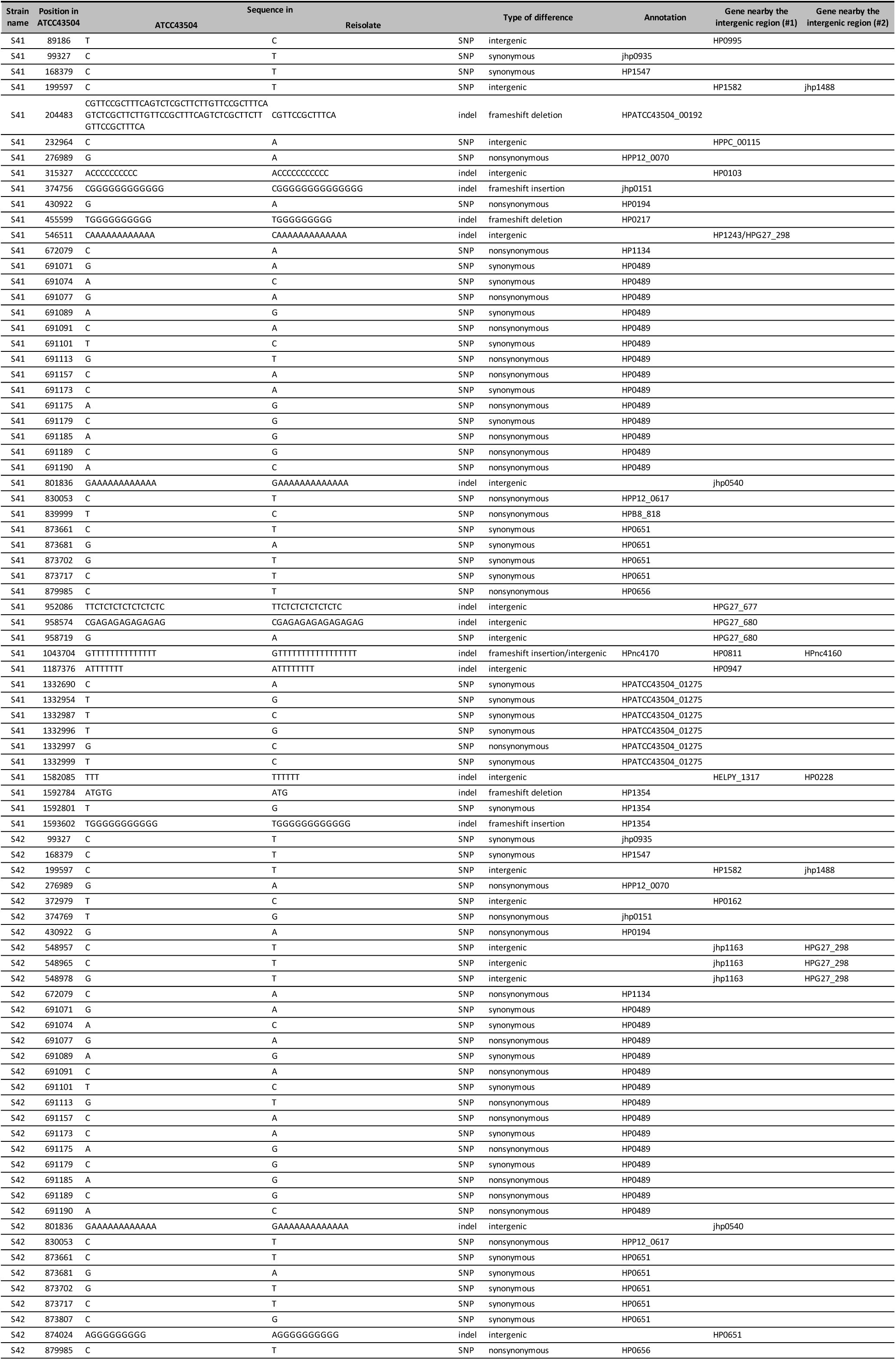
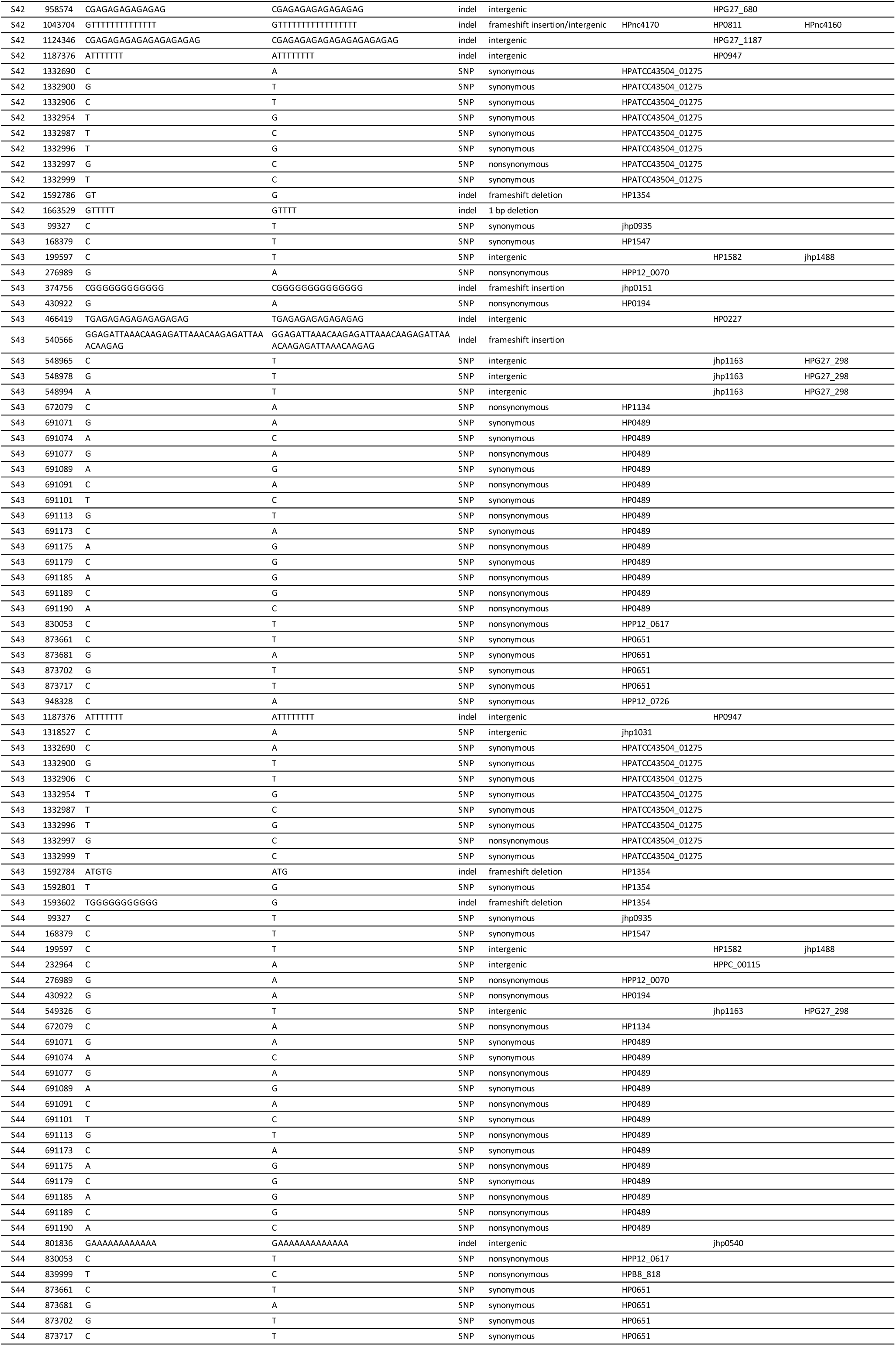
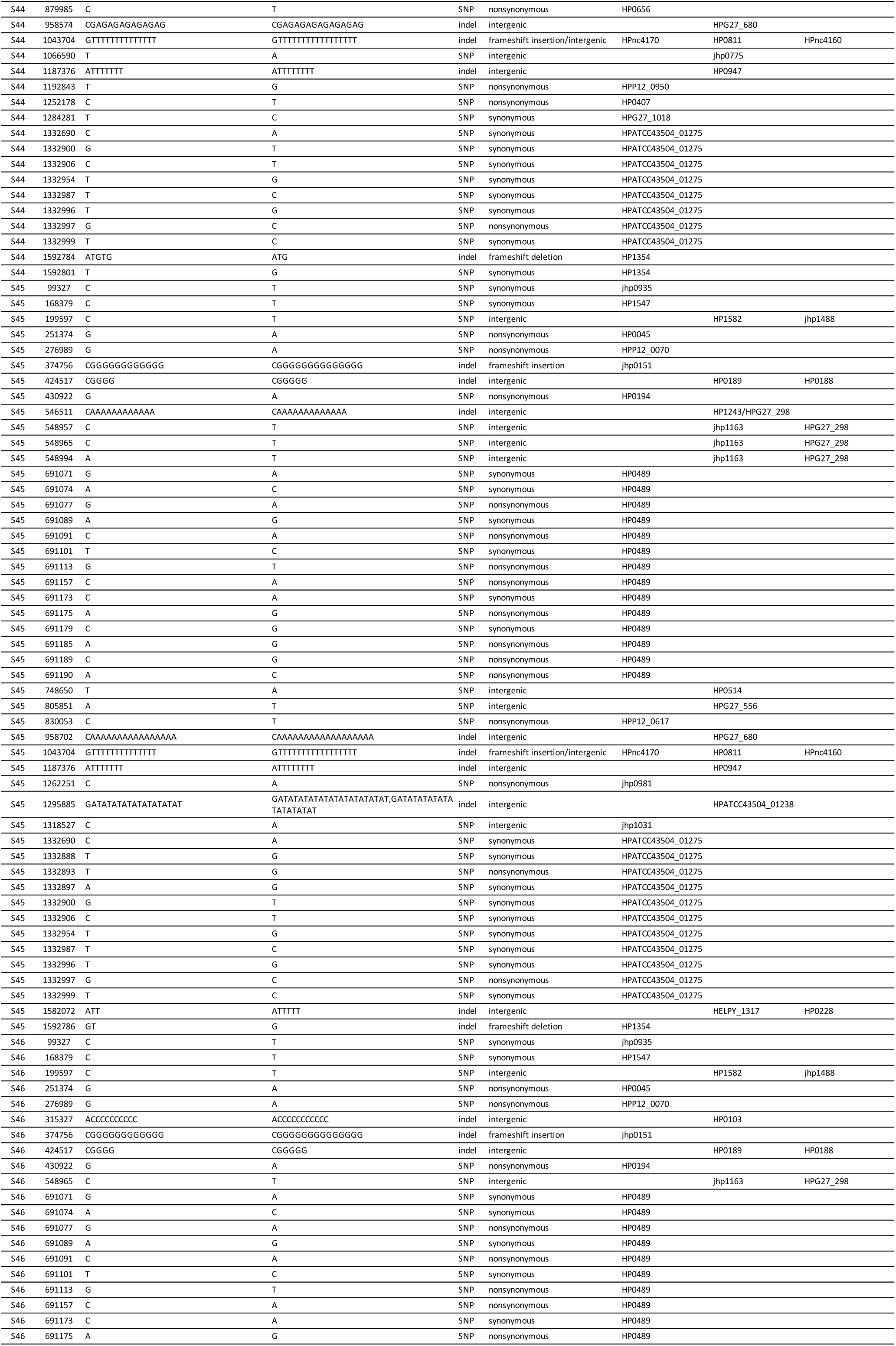
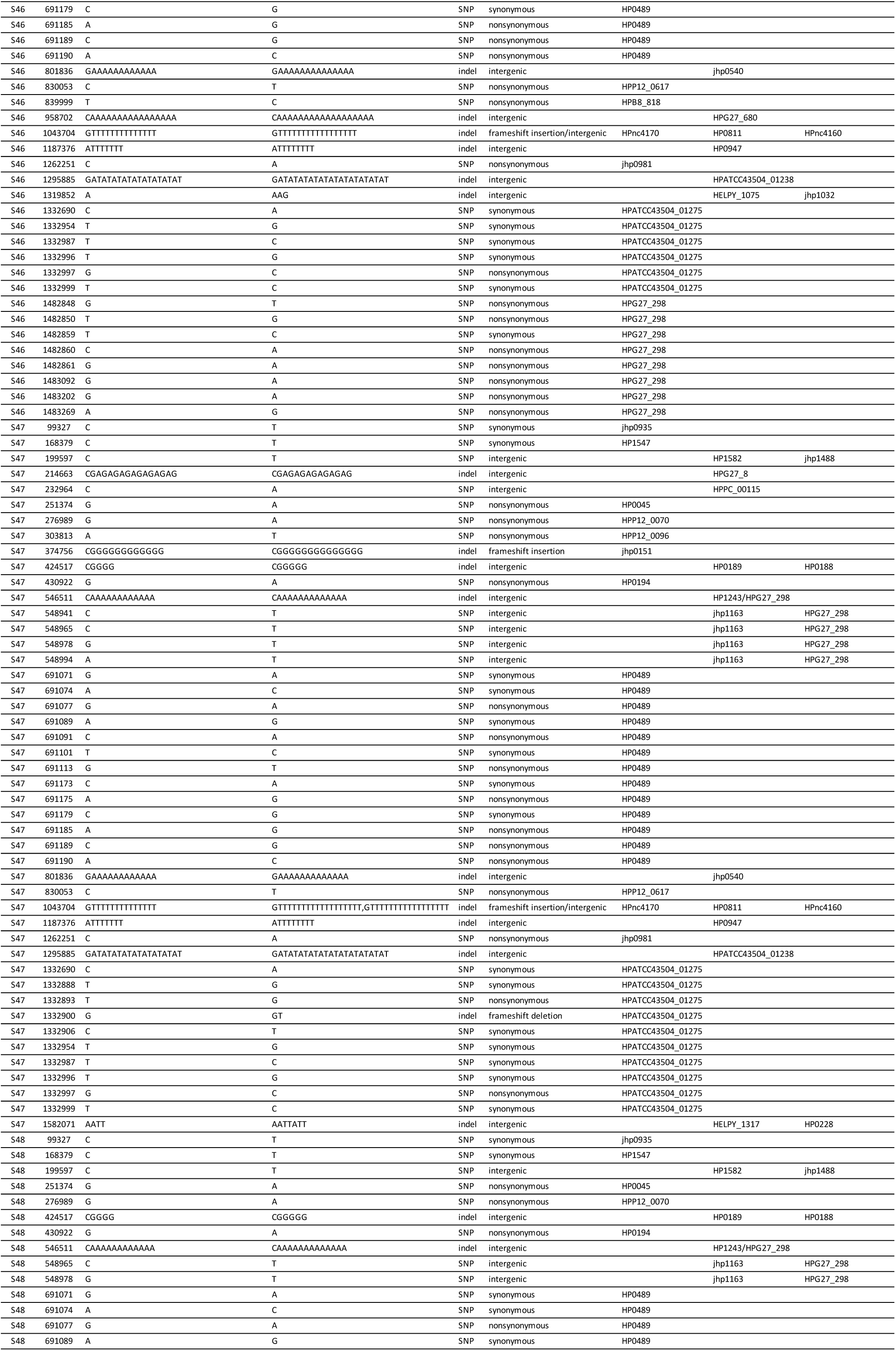
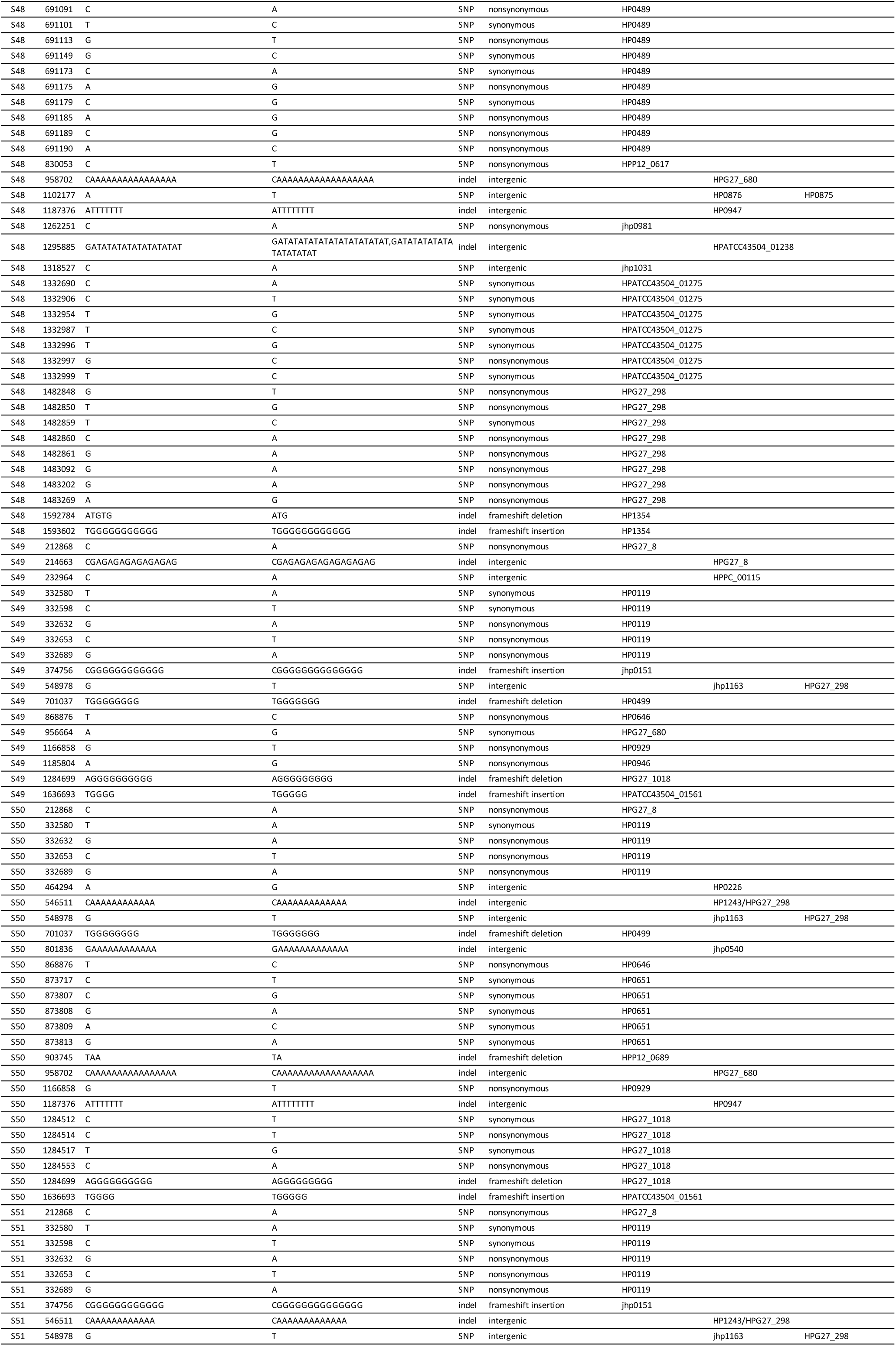
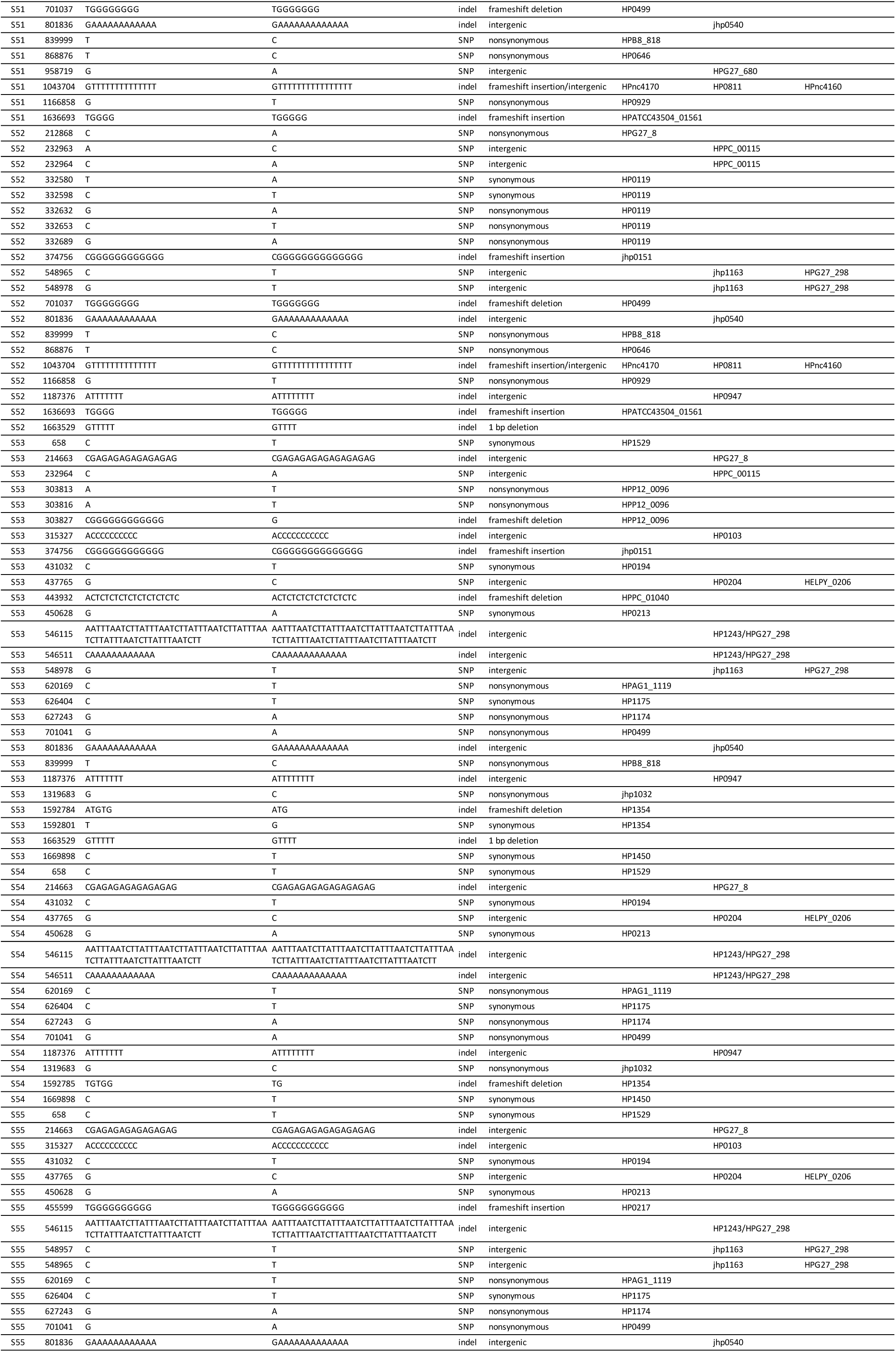
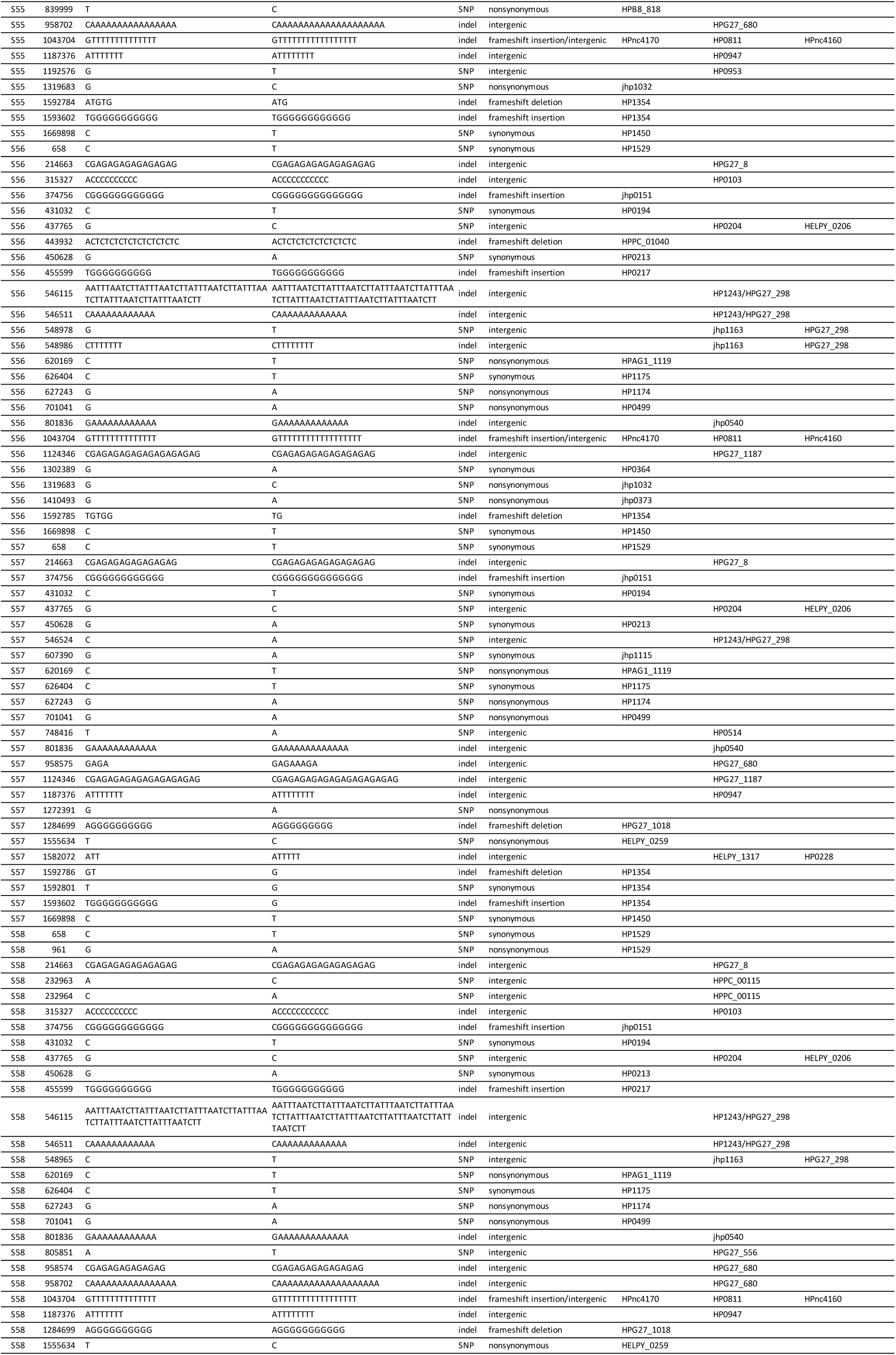
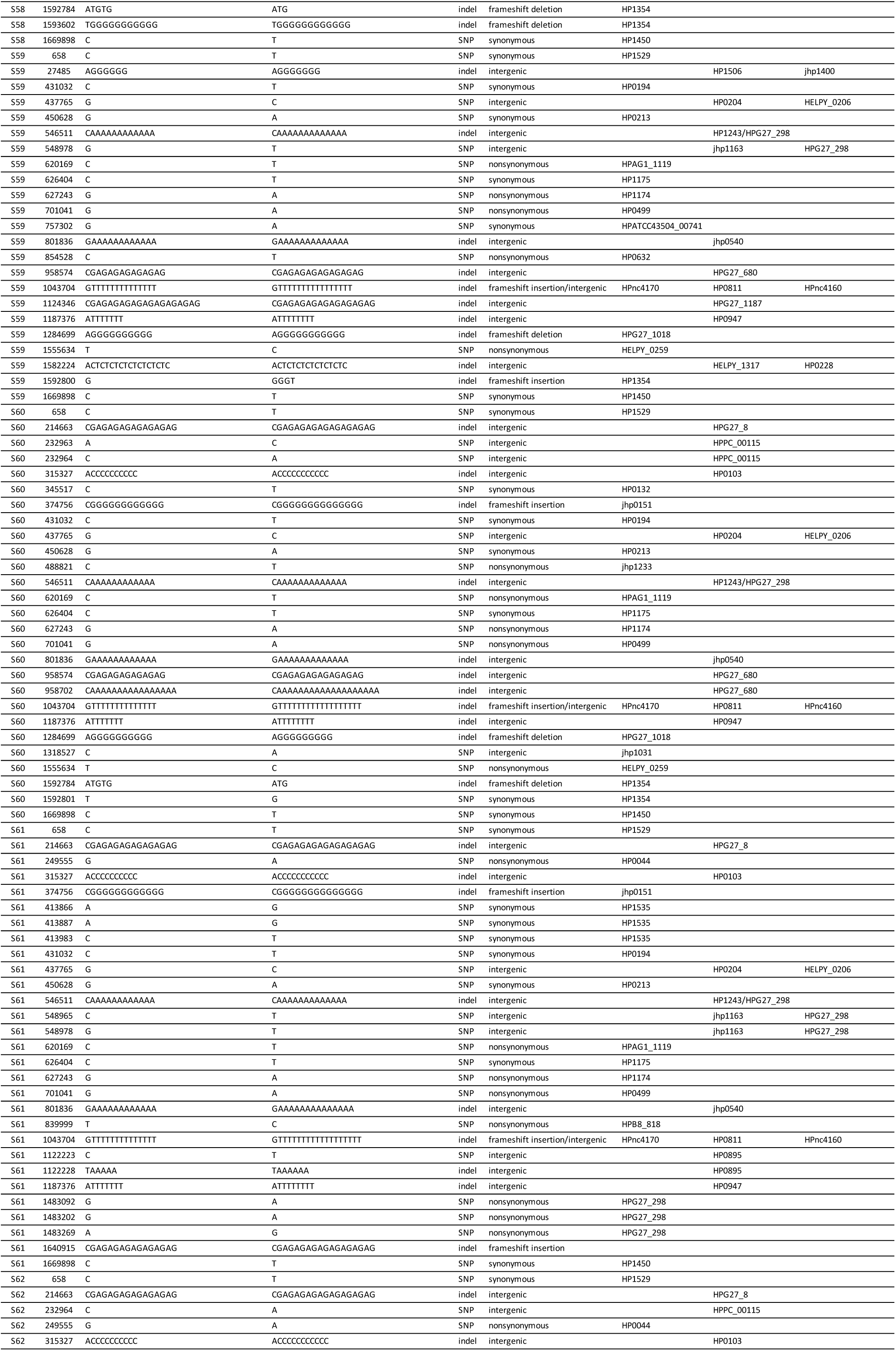
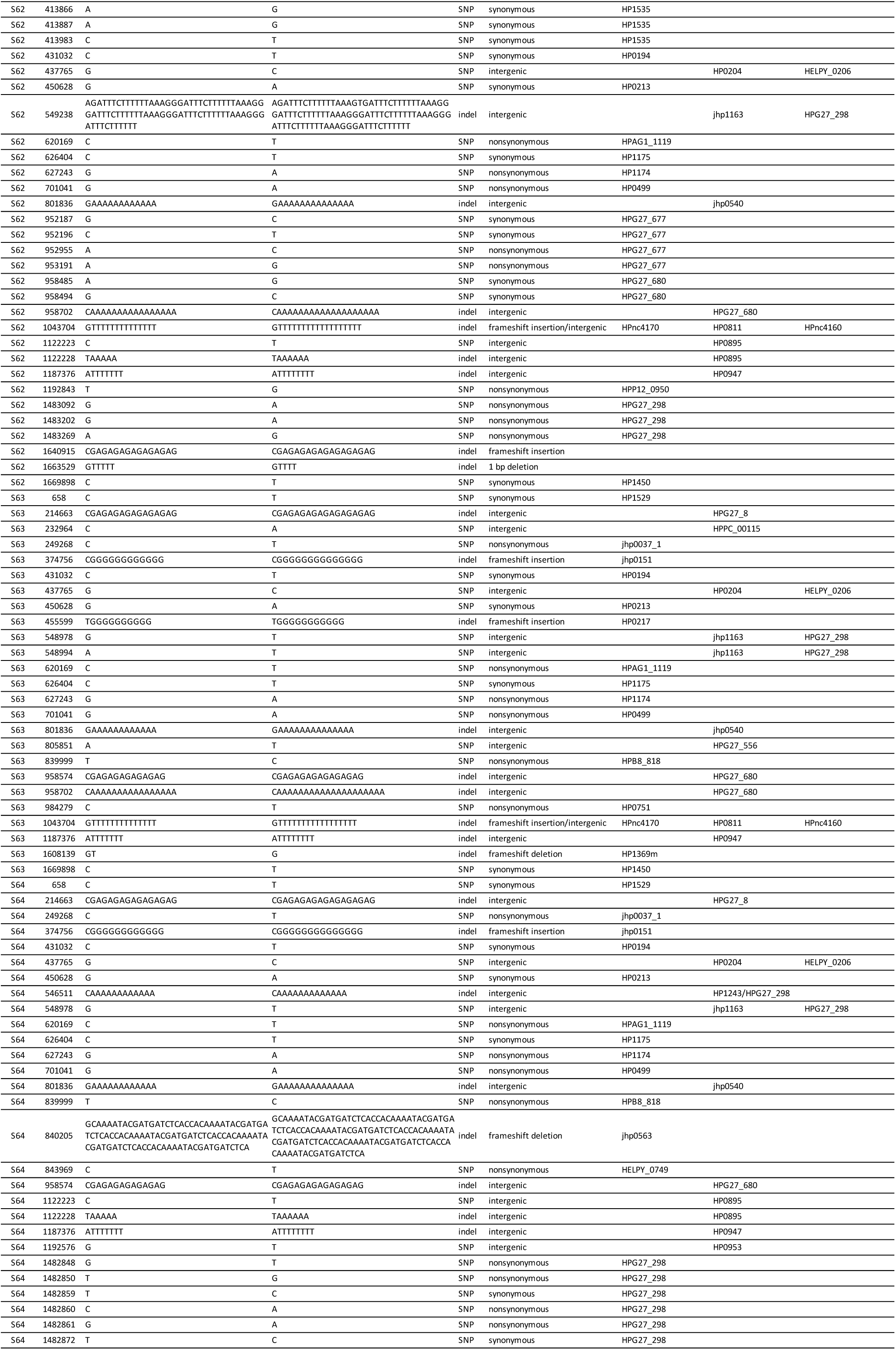
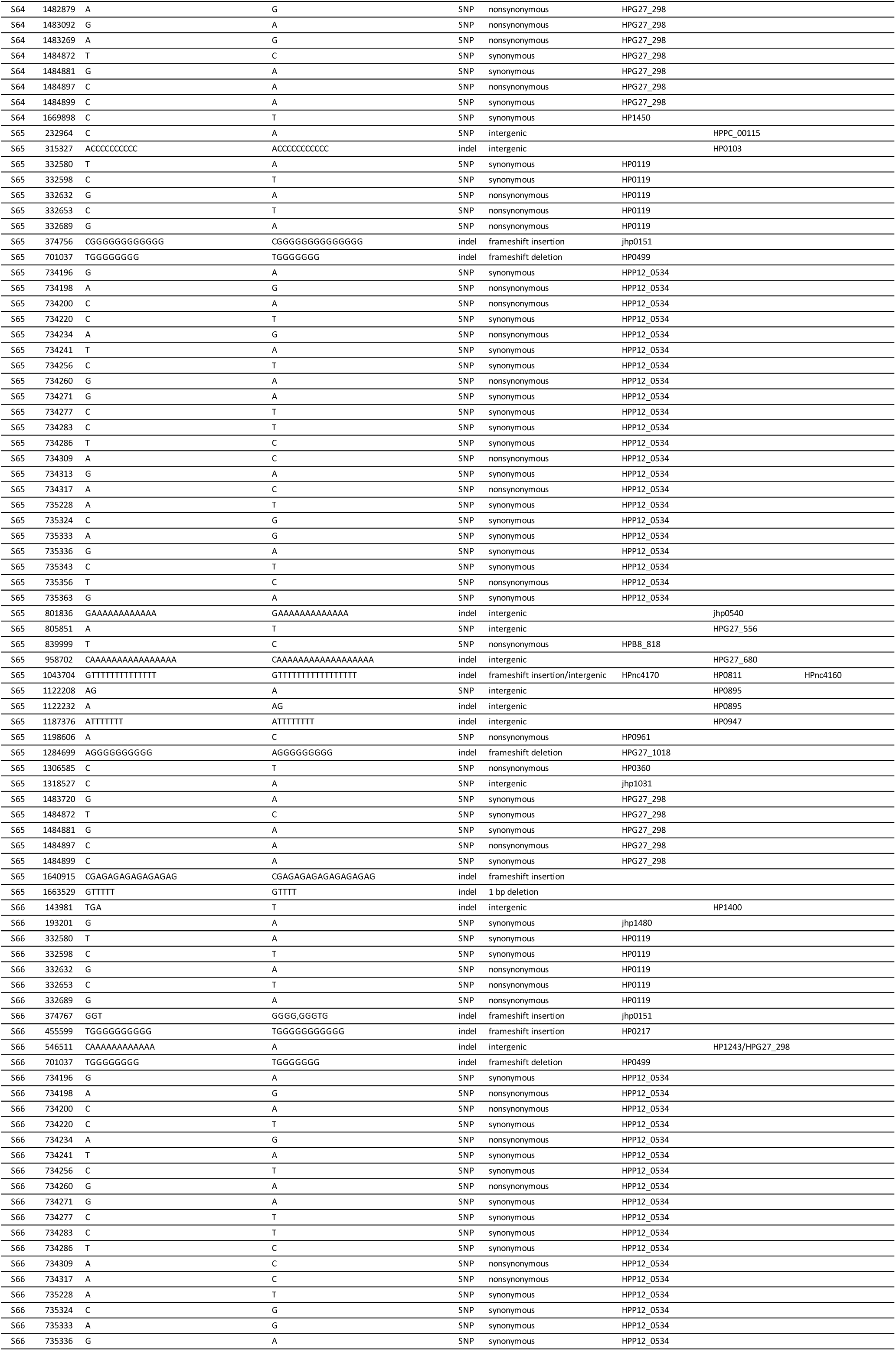
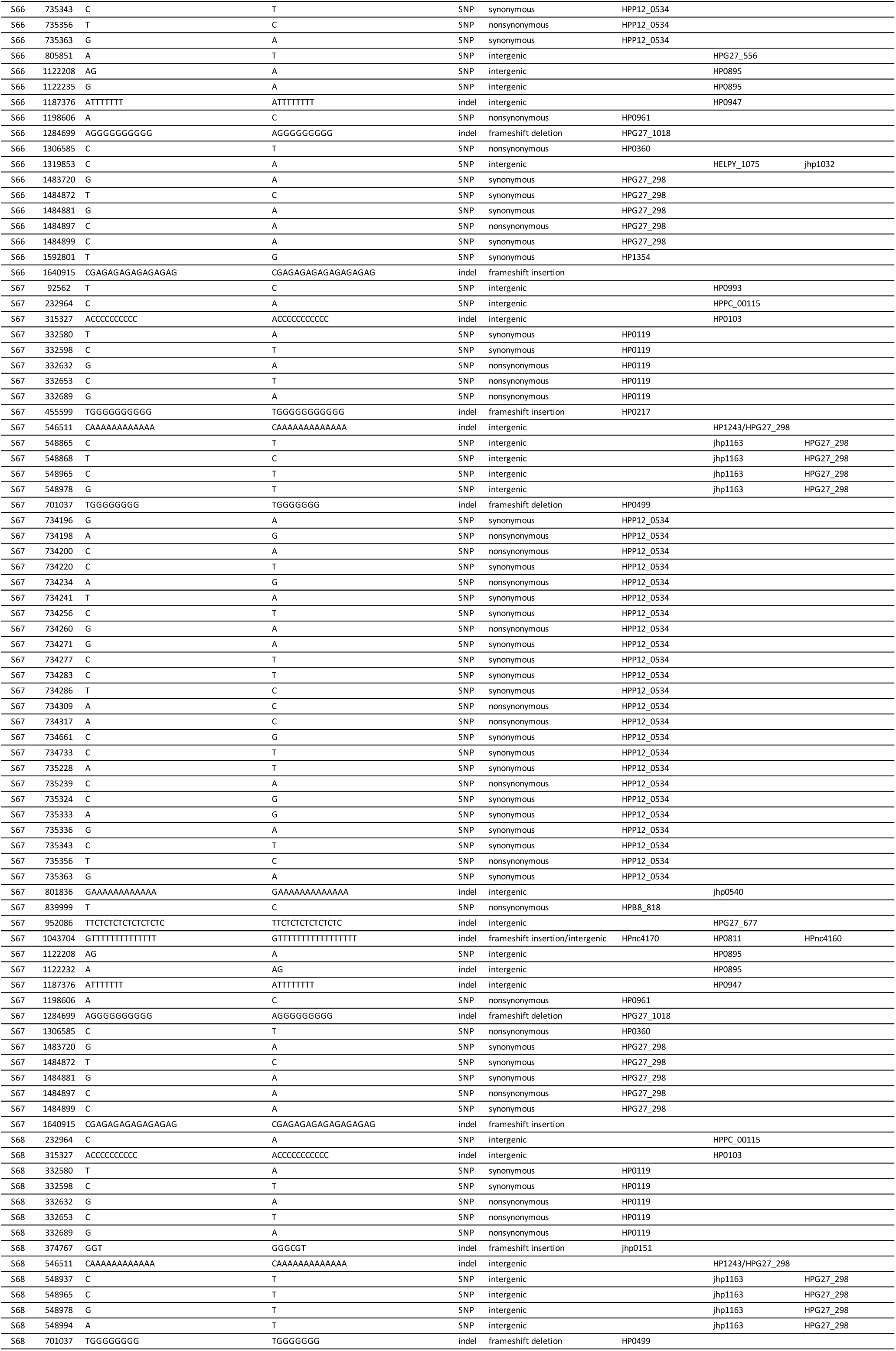
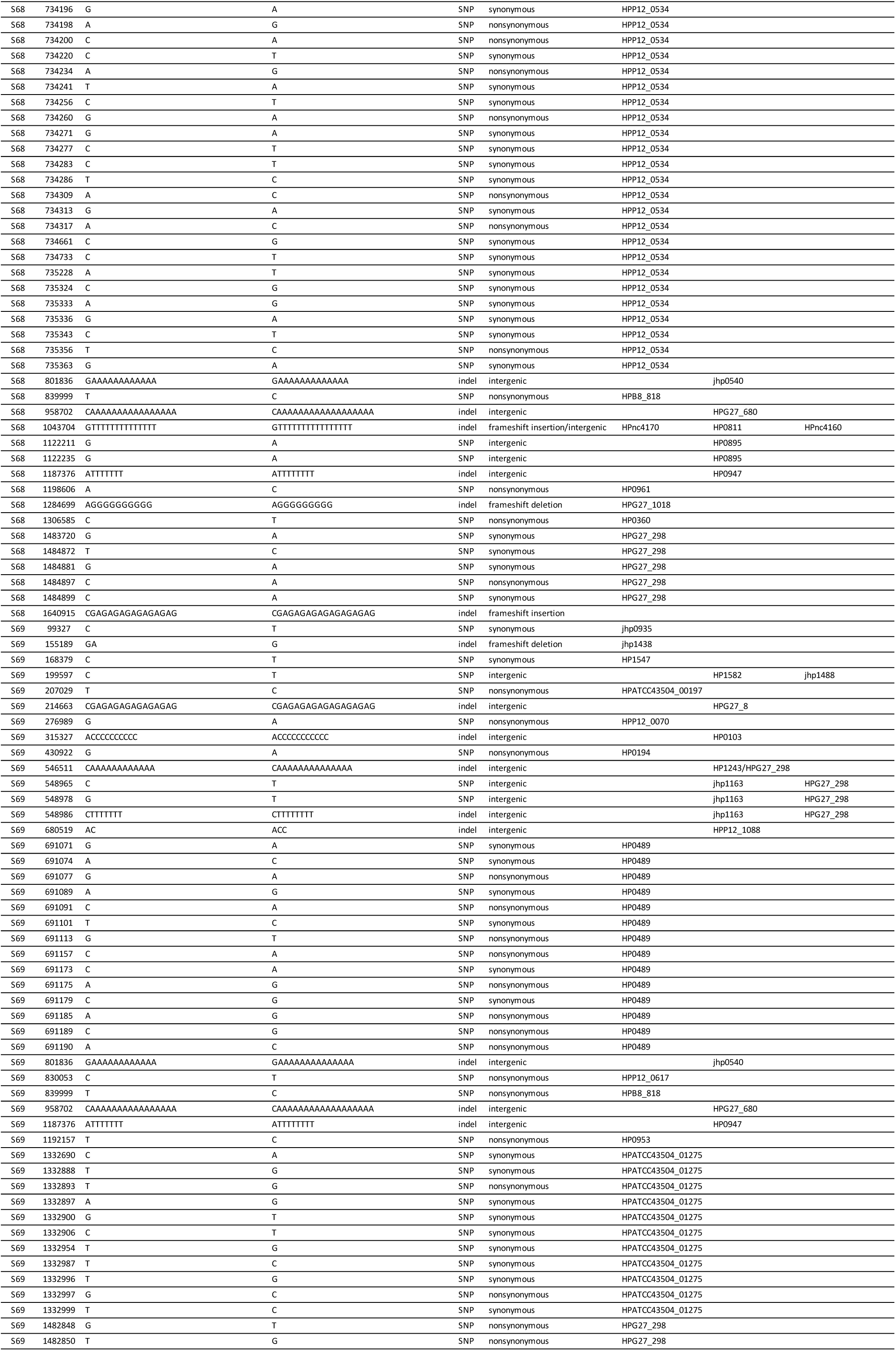
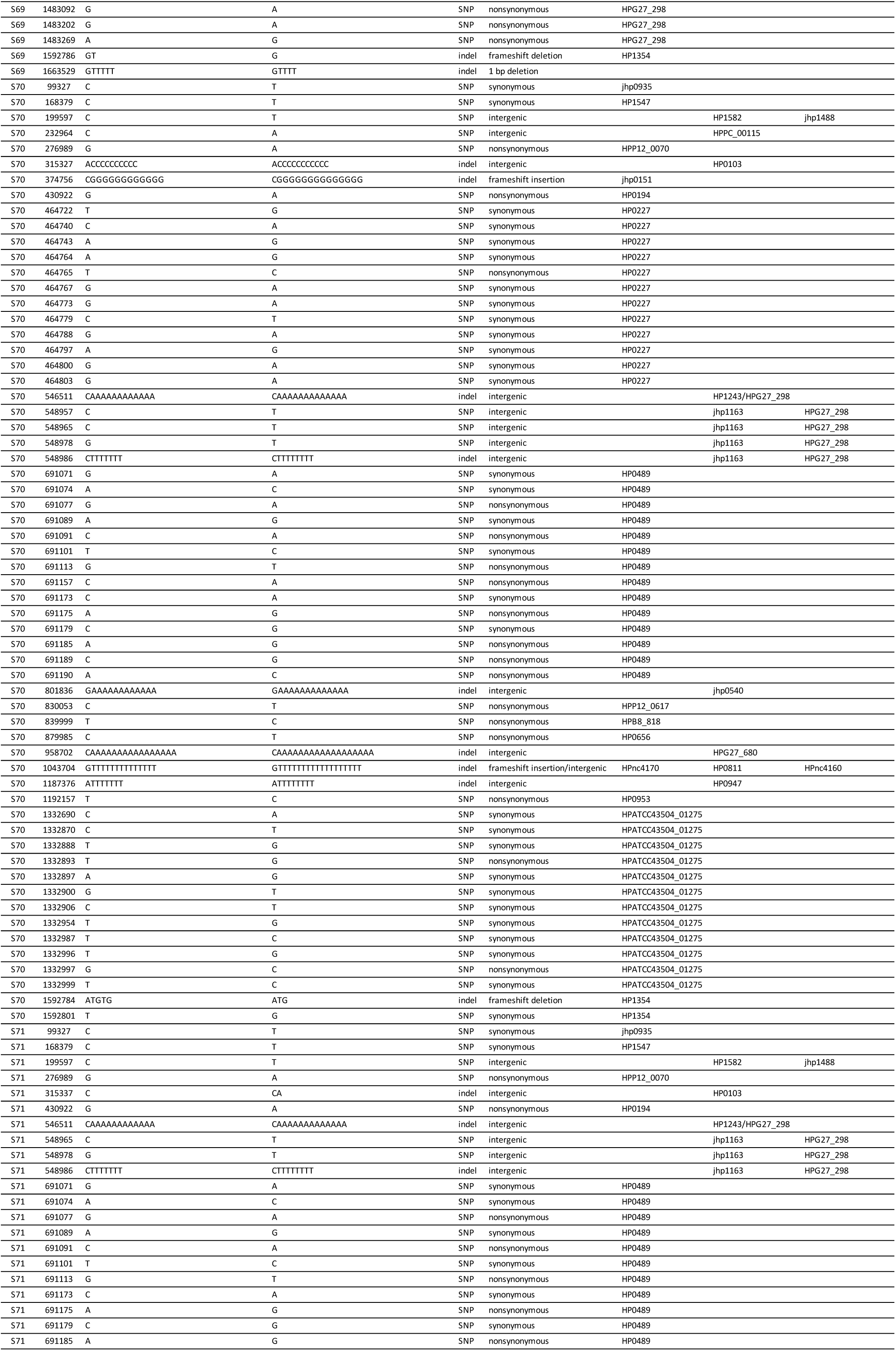
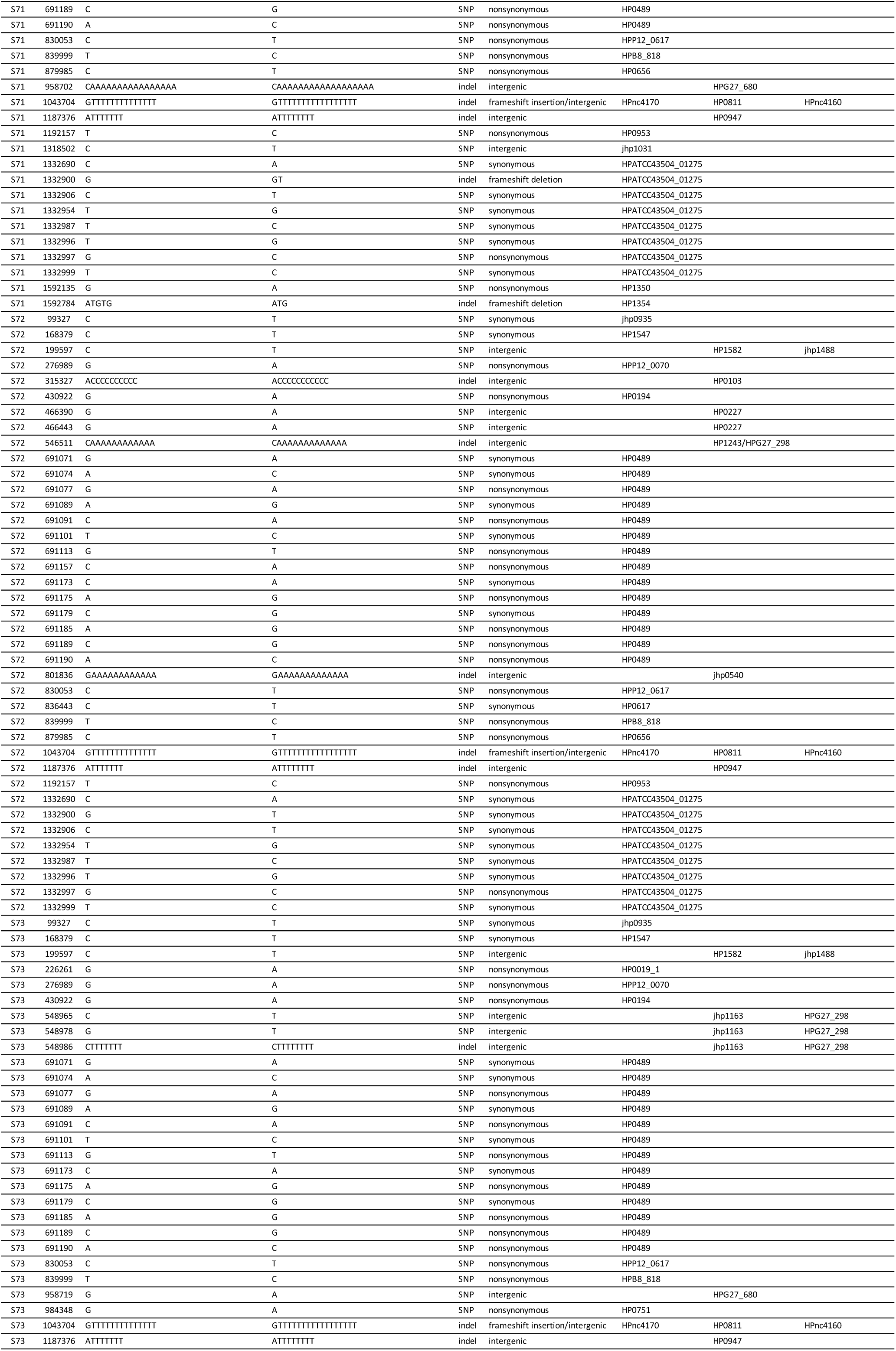
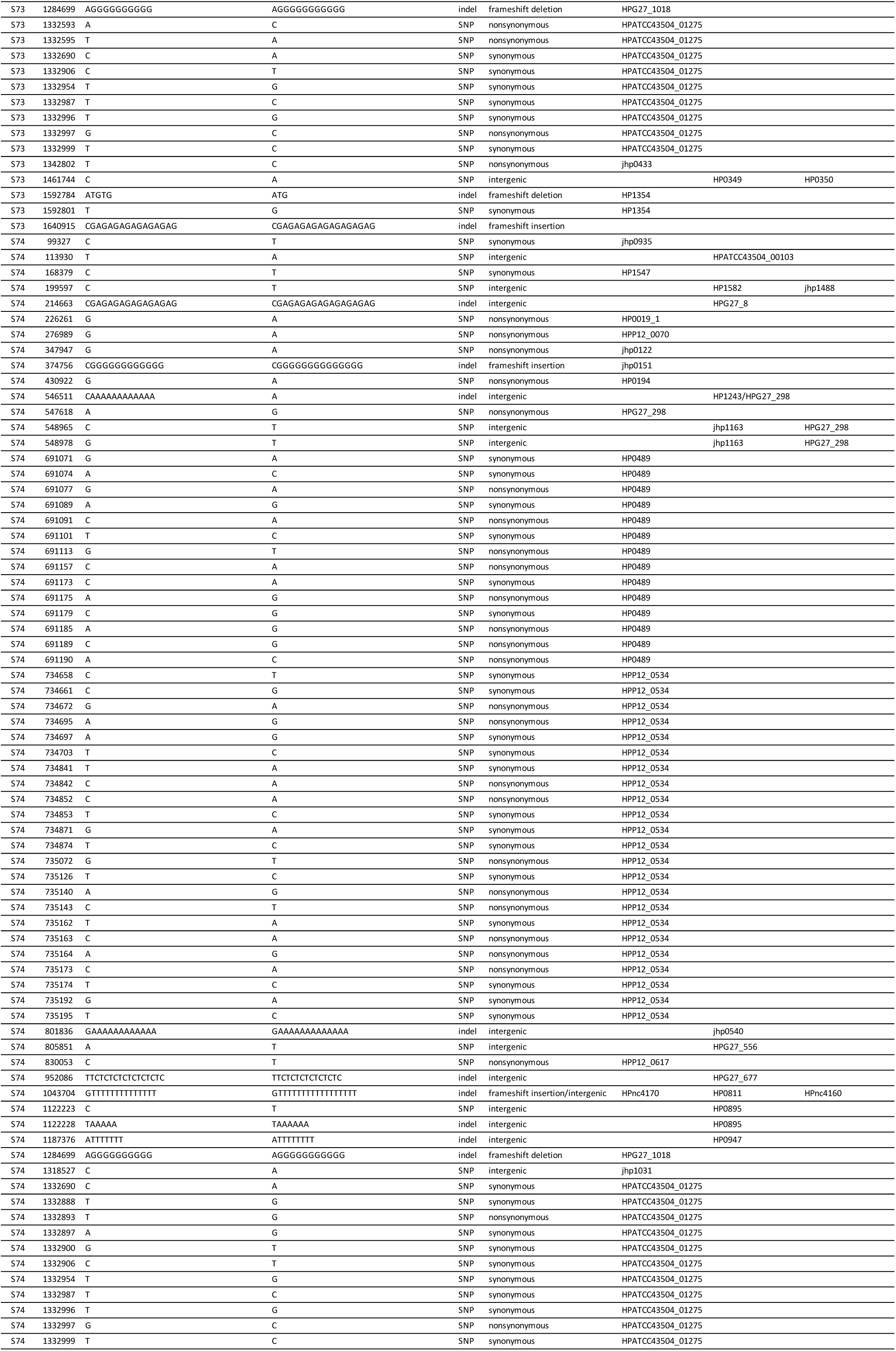
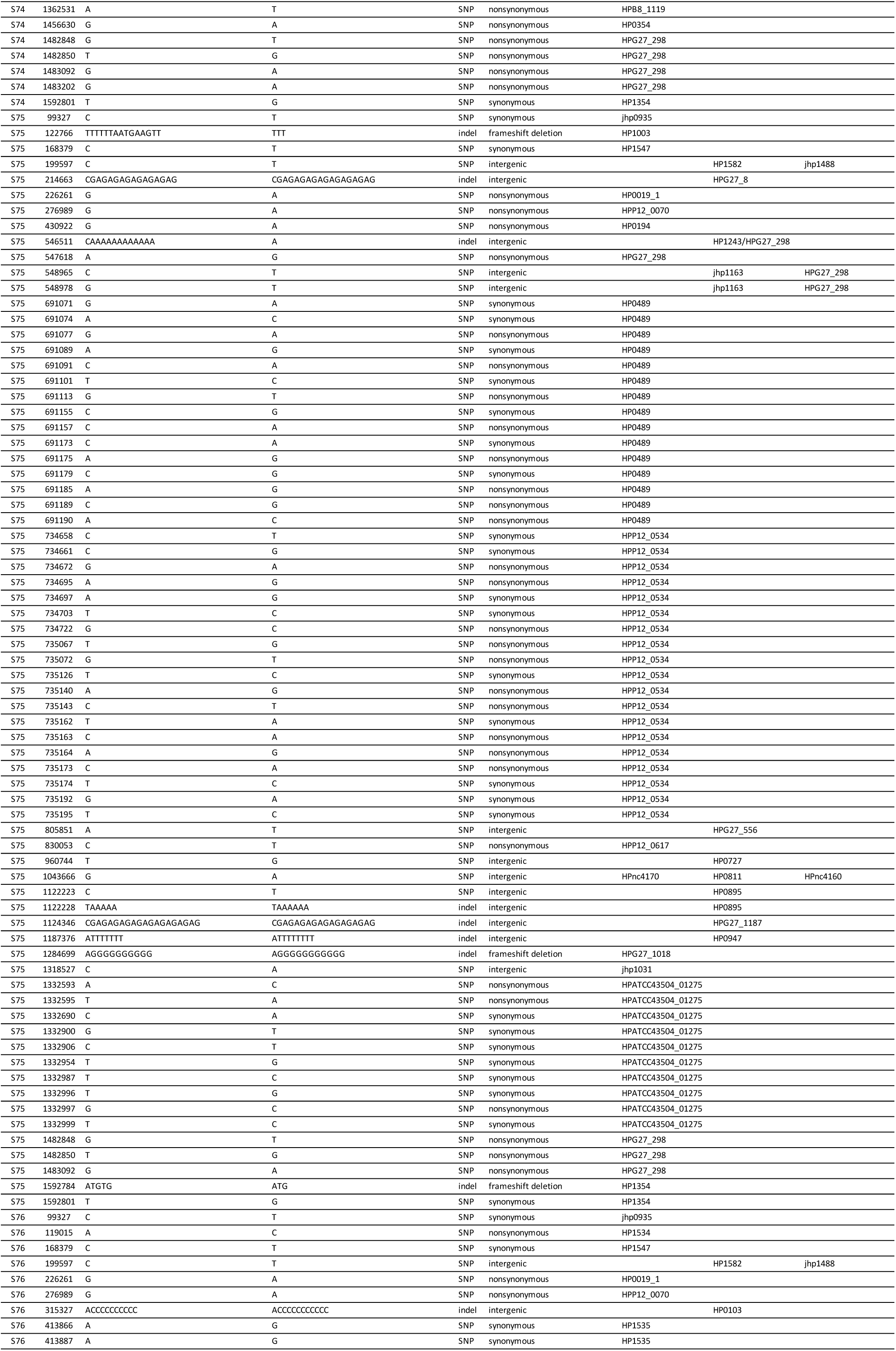
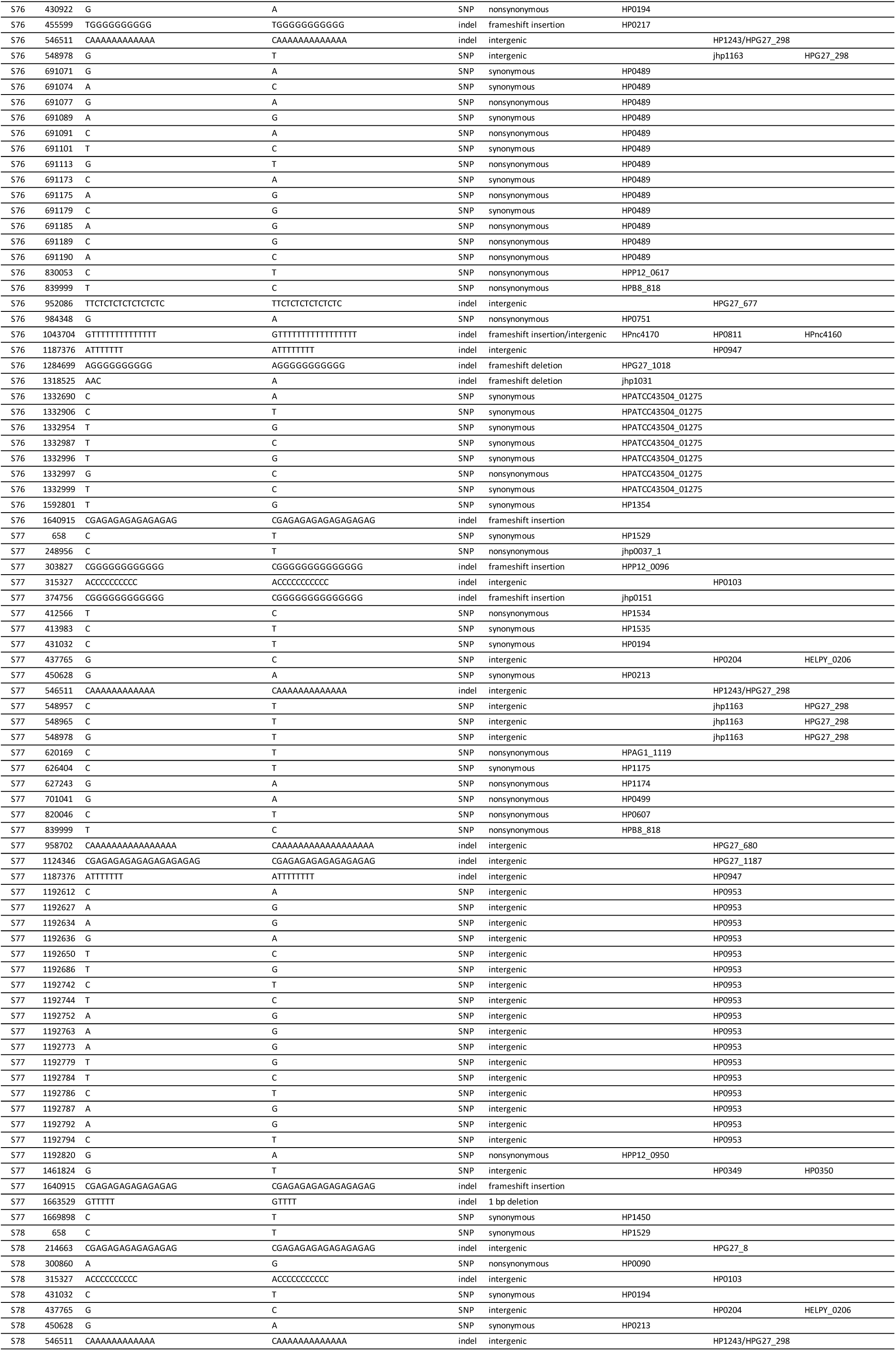
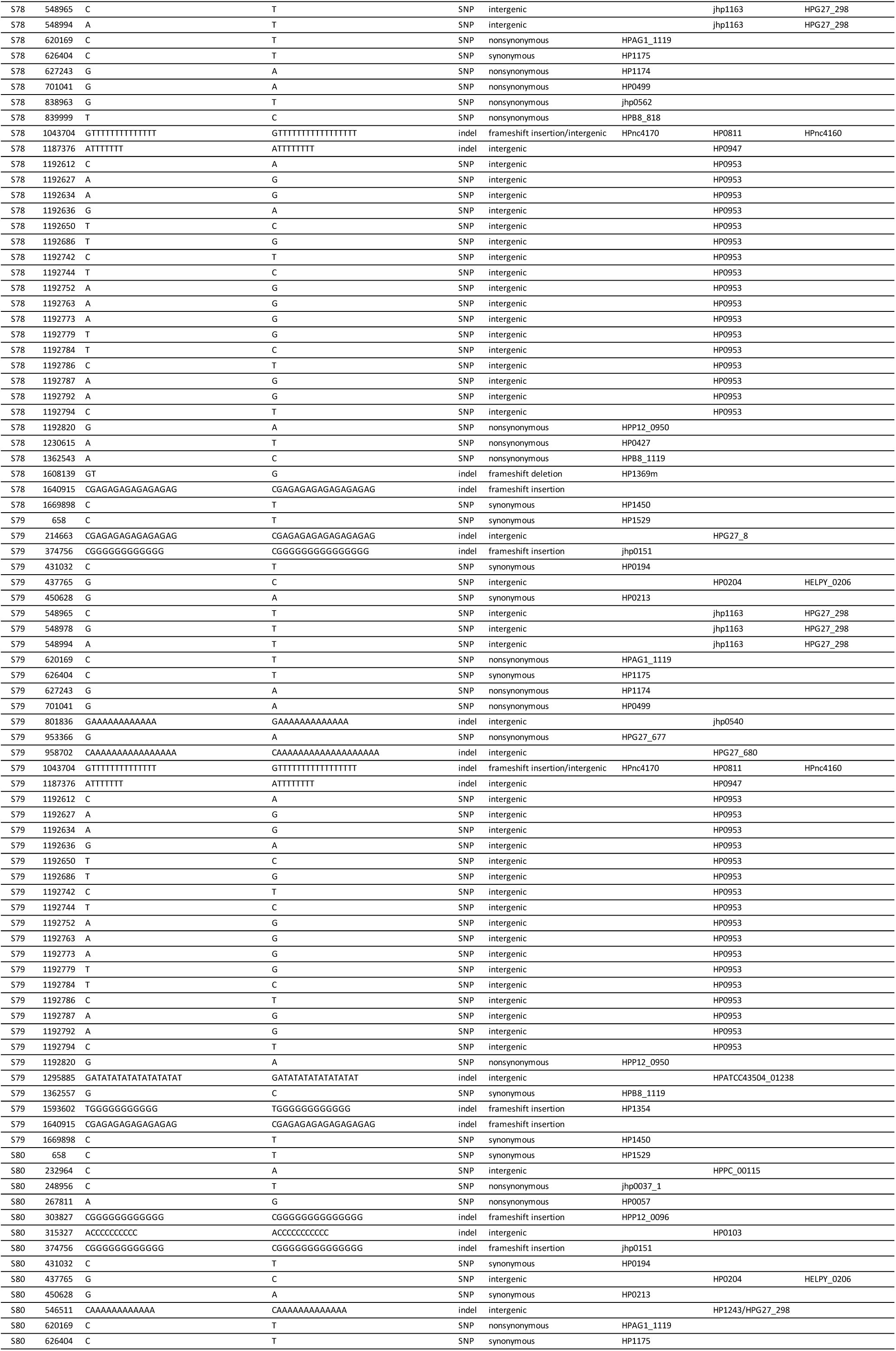
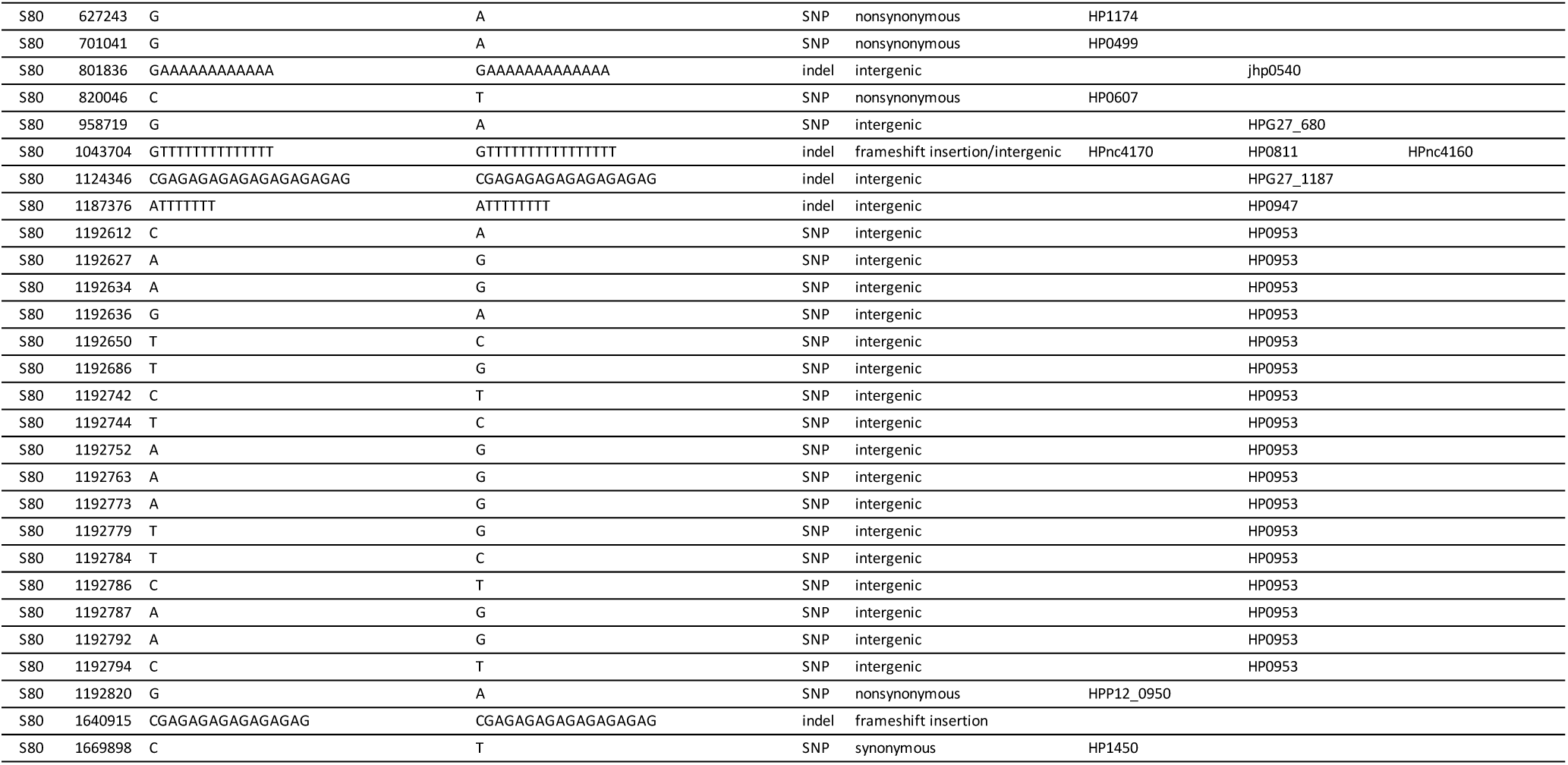
The mutated sequence list of 40 strains recovered from *H. pylori*-infected Mongolian gerbils’ stomach.

**Supplementary information 3.** Summary of mutations in the isolates recovered from *H. pylori*-infected C57BL/6 mice. The number of mutations in the isolates of 10 strains recovered from H. pylori-infected mice stomachs 8 weeks after post-infection were listed.

| Animal No. | Strain Name | Total No. of mutations | SNPs (single nucleotide polymorphysms) |  |  |  | indel (insertion/deletion) |  |  | SNPs rate per base per year | indel rate per base per year |
| --- | --- | --- | --- | --- | --- | --- | --- | --- | --- | --- | --- |
|  |  |  | total | nonsynon ymous | synonymo us | intergenic | Total | genic | intergenic |  |  |
| #1 | Hp1 | 37 | 13 | 5 | 7 | 1 | 24 | 8 | 16 | 5.04E-05 | 9.31E-05 |
|  | Hp2 | 36 | 16 | 5 | 9 | 2 | 20 | 7 | 13 | 6.20E-05 | 7.76E-05 |
| #2 | Hp3 | 38 | 16 | 6 | 8 | 2 | 22 | 8 | 14 | 6.20E-05 | 8.53E-05 |
|  | Hp4 | 38 | 18 | 6 | 9 | 3 | 20 | 7 | 13 | 6.98E-05 | 7.76E-05 |
| #3 | Hp5 | 34 | 15 | 6 | 7 | 2 | 19 | 6 | 13 | 5.82E-05 | 7.37E-05 |
|  | Hp6 | 33 | 10 | 4 | 5 | 1 | 23 | 7 | 16 | 3.88E-05 | 8.92E-05 |
| #4 | Hp7 | 45 | 25 | 11 | 12 | 2 | 20 | 10 | 10 | 9.69E-05 | 7.76E-05 |
|  | Hp8 | 42 | 19 | 7 | 10 | 2 | 23 | 8 | 15 | 7.37E-05 | 8.92E-05 |
| #5 | Hp9 | 46 | 30 | 11 | 17 | 2 | 16 | 5 | 11 | 1.16E-04 | 6.20E-05 |
|  | Hp10 | 36 | 13 | 5 | 7 | 1 | 23 | 8 | 15 | 5.04E-05 | 8.92E-05 |
| Average |  |  |  |  |  |  |  |  |  | 6.79E-05 | 8.14E-05 |
| SD |  |  |  |  |  |  |  |  |  | 2.32E-05 | 9.50E-06 |

**Supplementary information 4.**
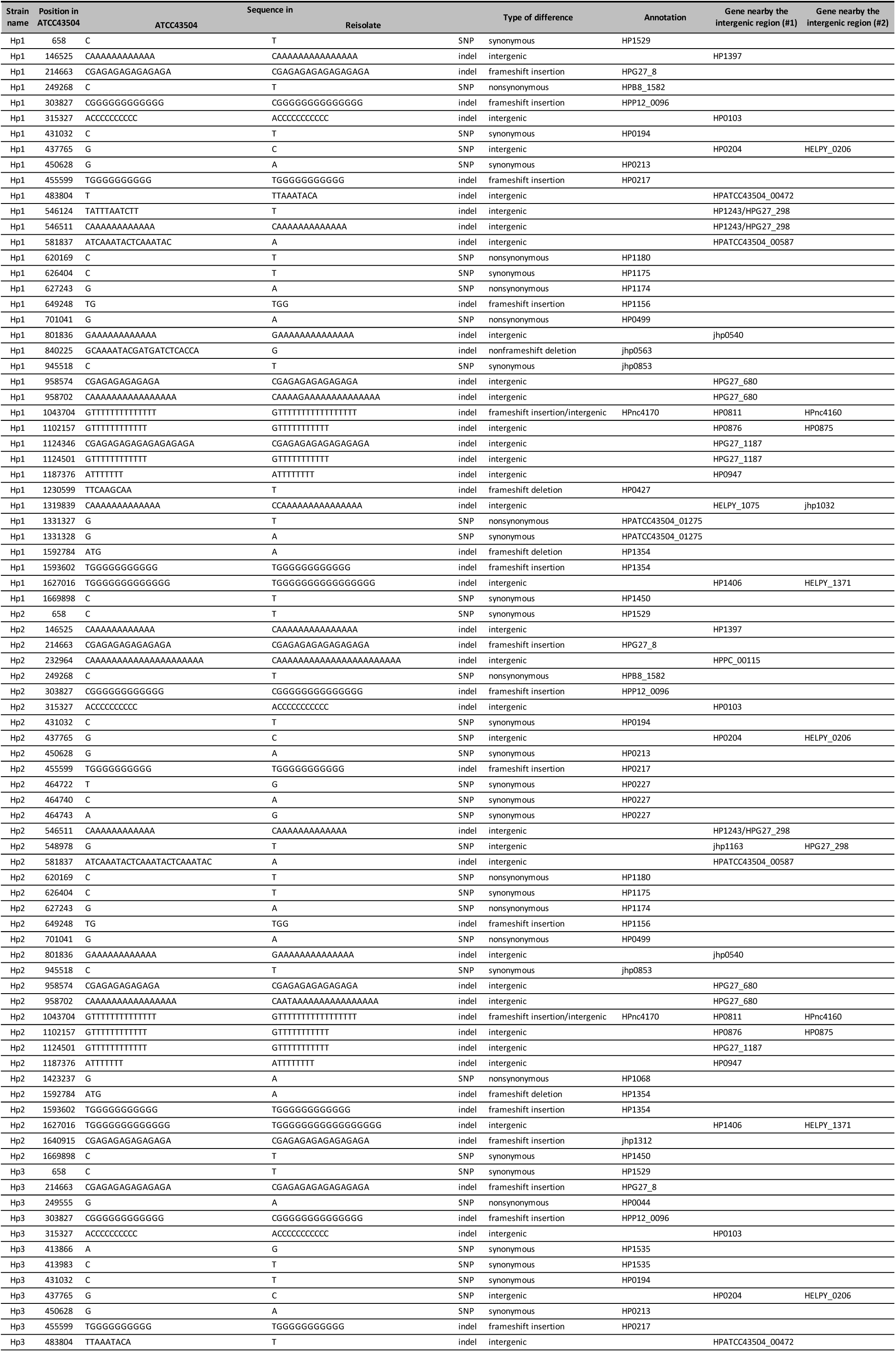
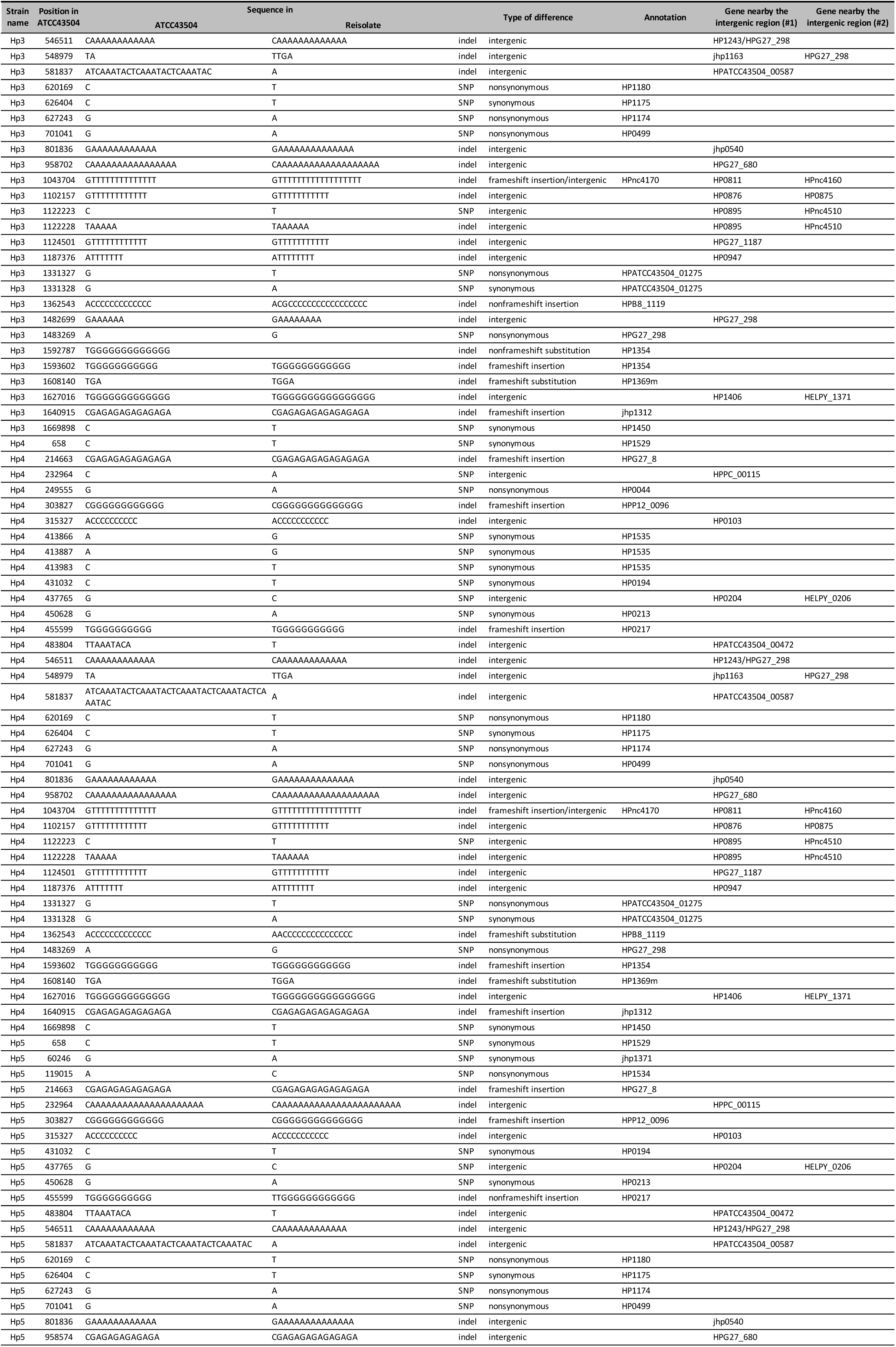
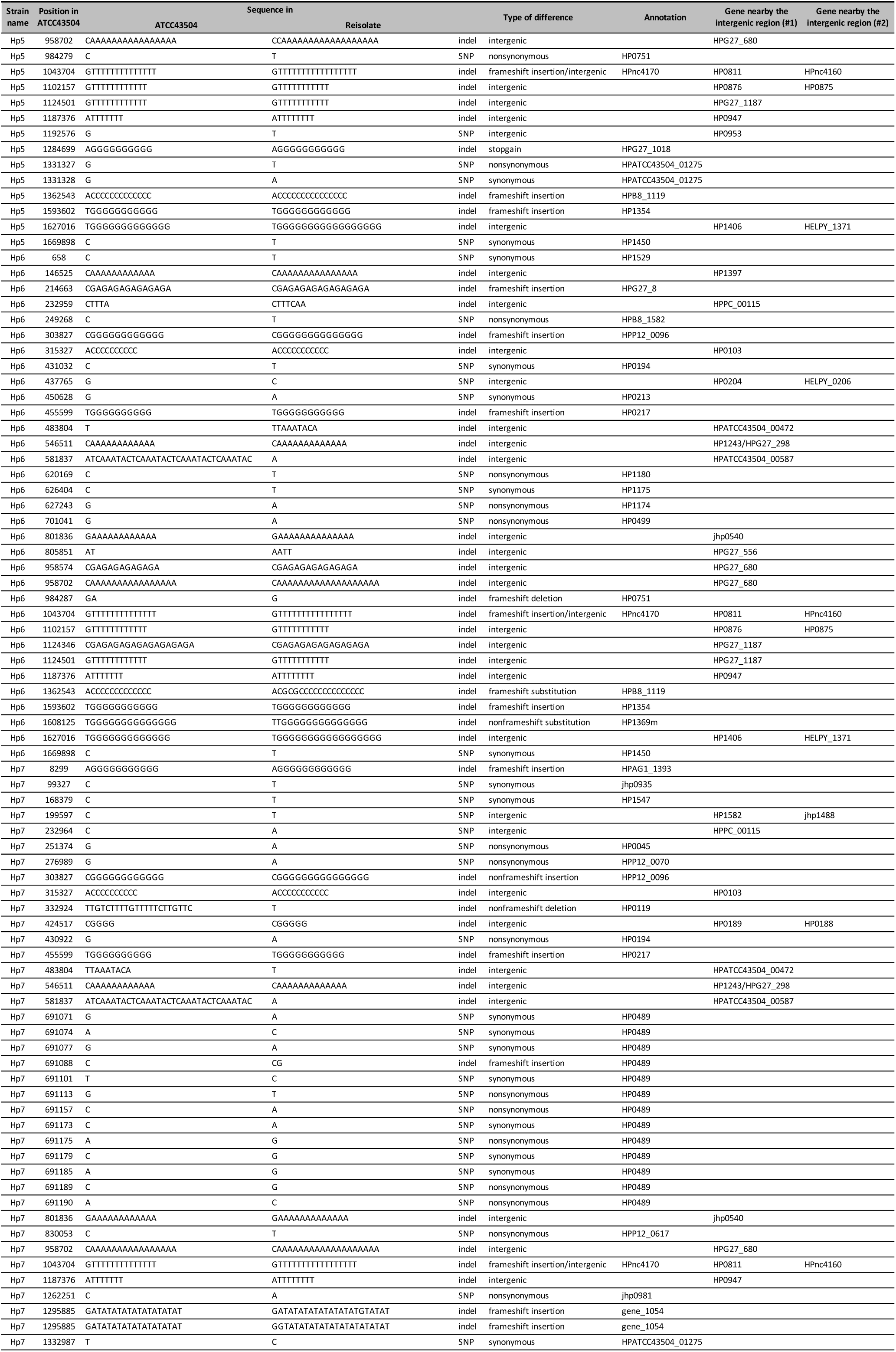
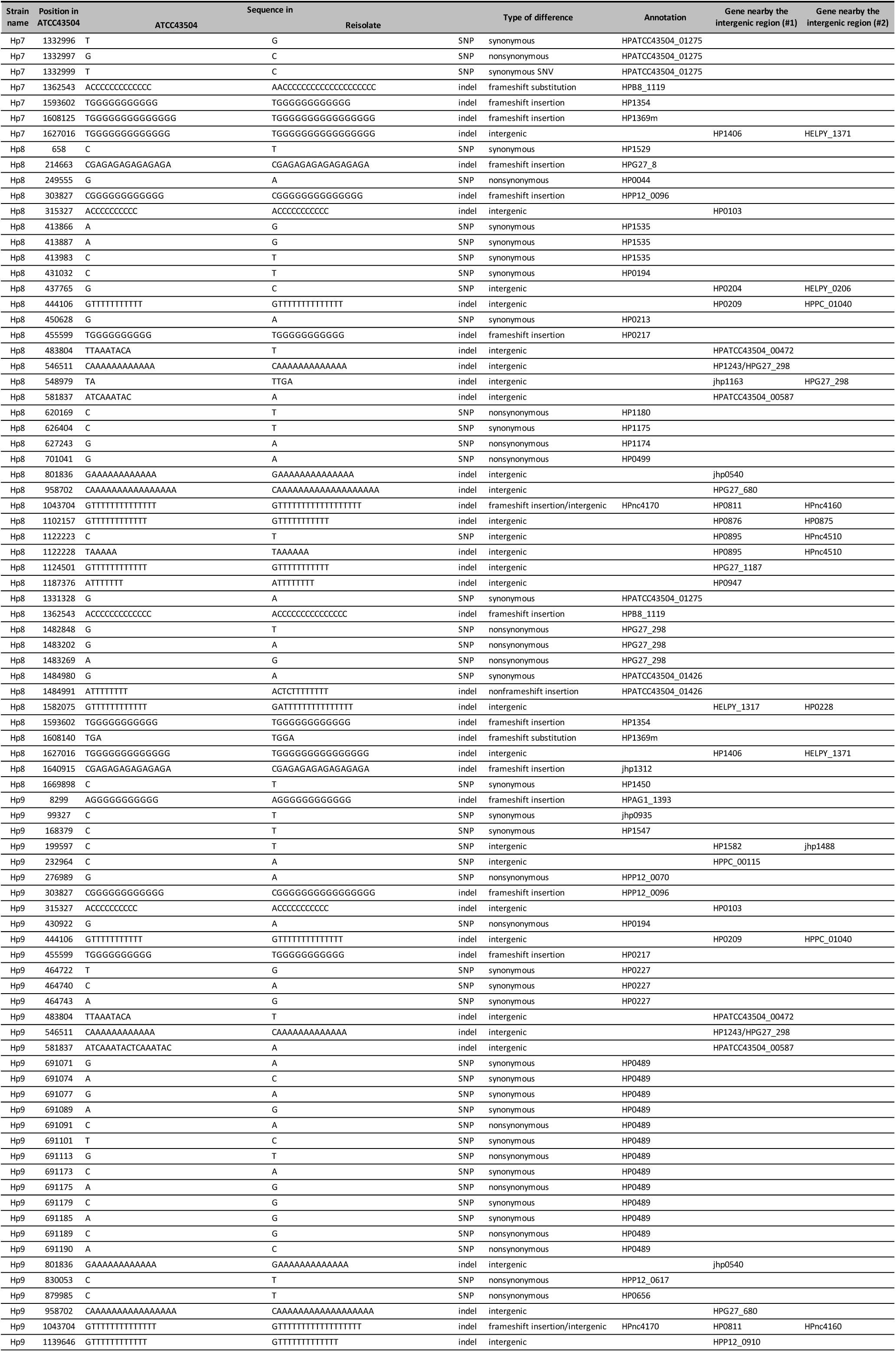
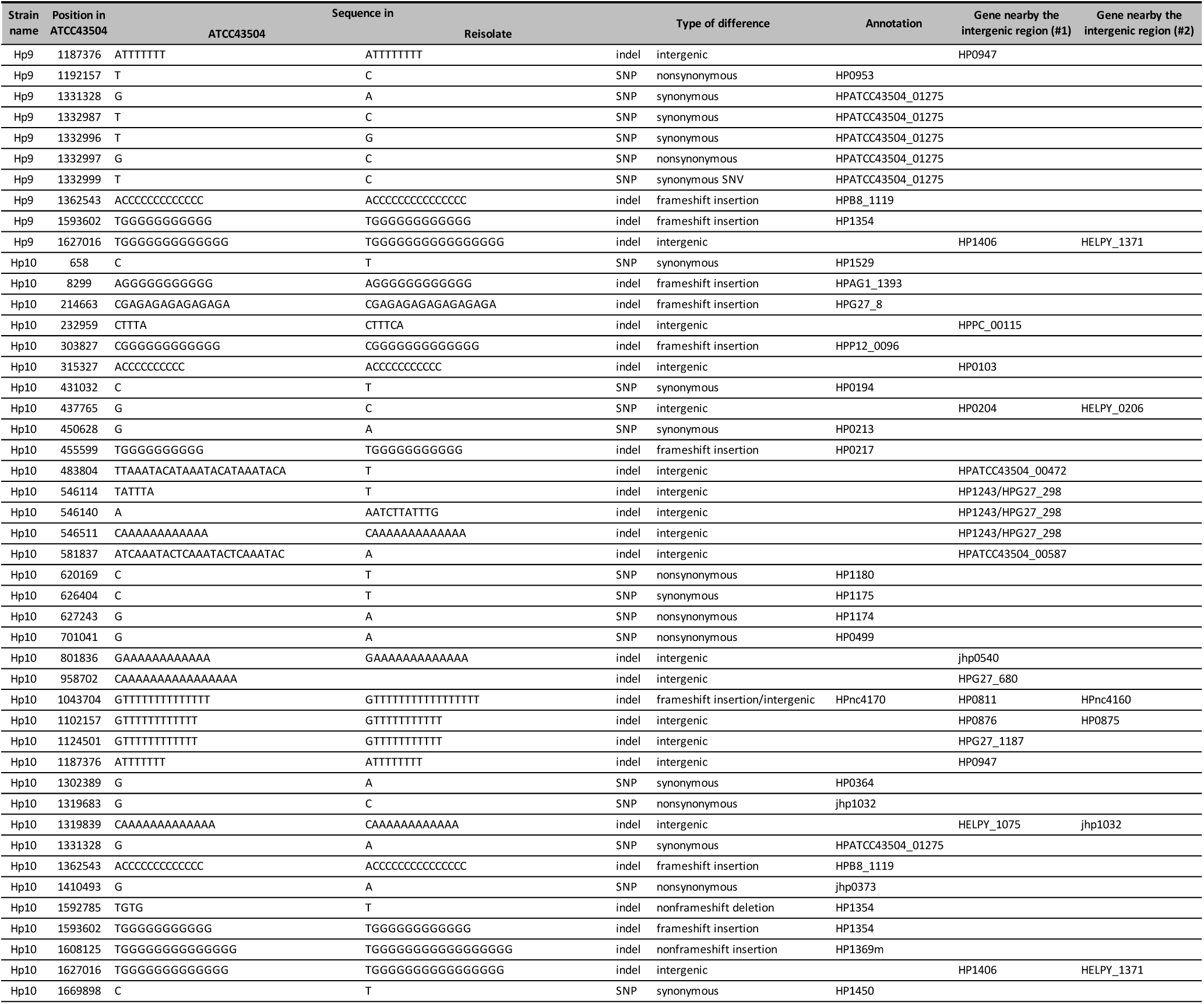
The mutated sequence list of 10 strains recovered from *H. pylori*-infected C57BL/6 mice stomach.

**Supplementary information 5.**
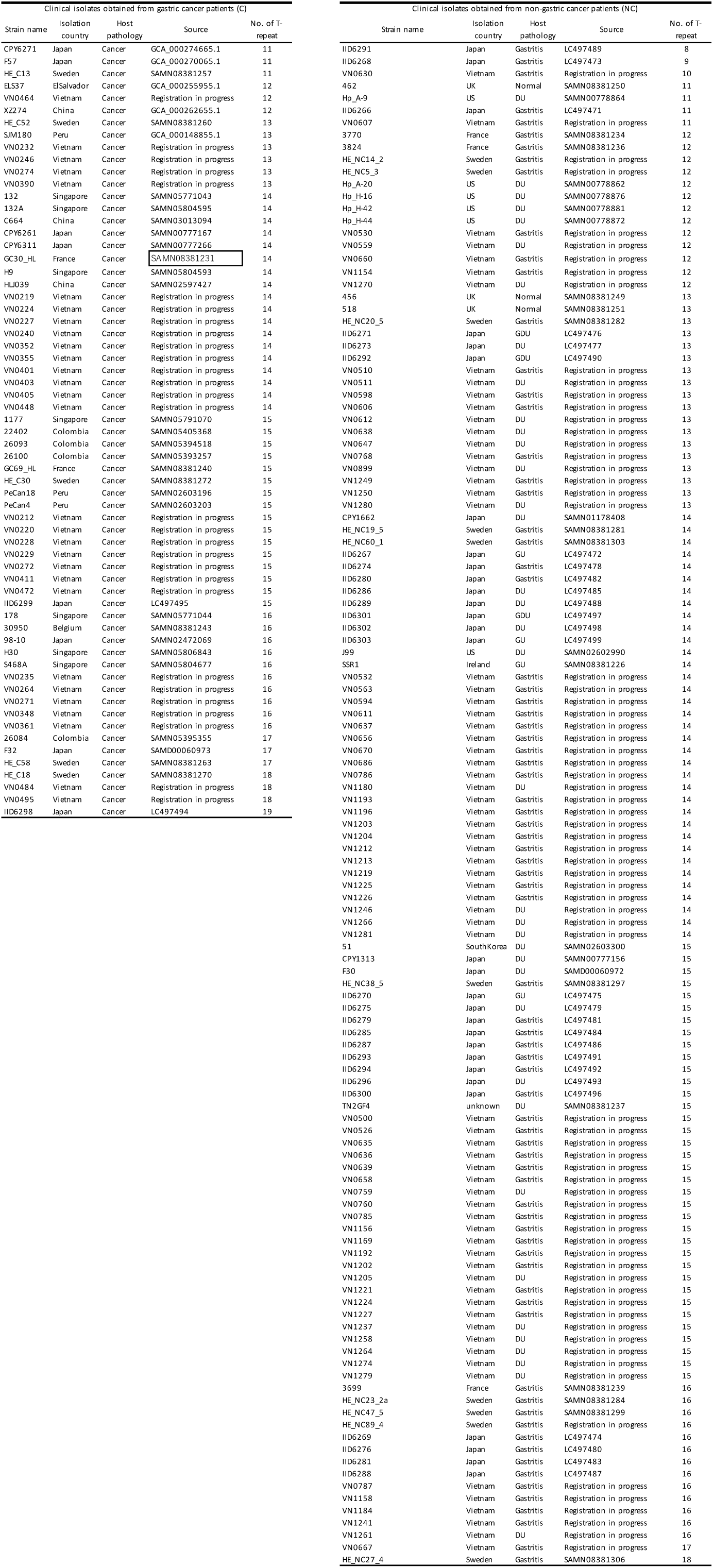
Information of *H. pylori* clinical isolates used in Fig. 1f, and Fig 4g.

**Supplementary information 6.**
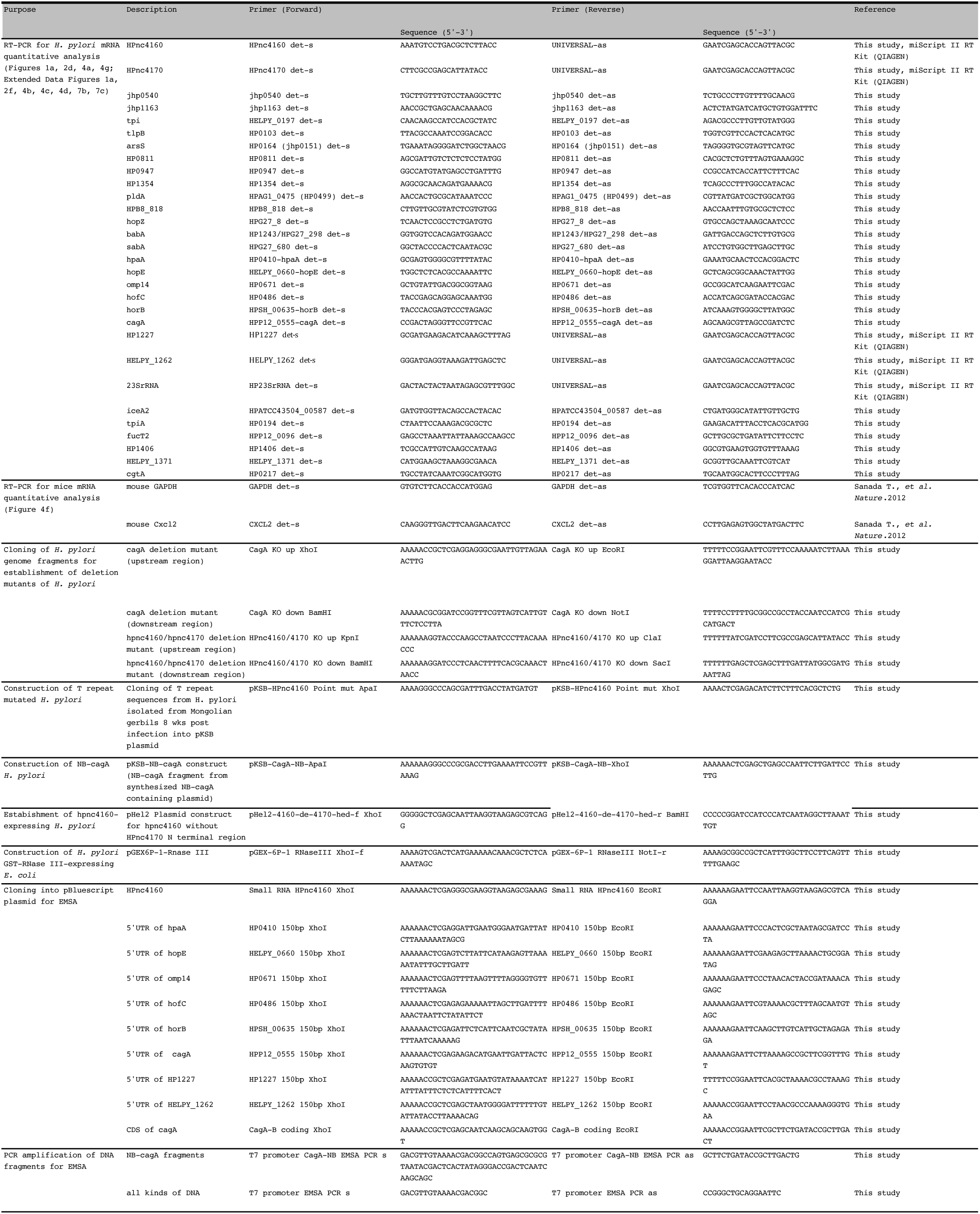
Primers used in this study.

## Acknowledgements

The authors would like to thank Manuel Amieva for providing us *H. pylori* strains. We gratefully acknowledge Keisuke Katsura for his support. We would like to thank the members of the Division of Bacteriology, Department of Infectious Diseases Control, International Research Center for Infectious Diseases, The Institute of Medical Science, The University of Tokyo, and the members of Department of Infection Microbiology, Research Institute for Microbial Diseases, Osaka University. This work was supported in part by Grant-in-Aid for Scientific Research from the Ministry of Education, Culture, Sports, Science, and Technology of Japan [17K19551, 18K07127, 19K22704 (to H.M.), 16K07083 (to T.S.), 17K14974 (to M.T.)], the Naito Foundation, the Tokyo Biochemical Research Foundation (to H.M. and P.S.), and the Kao Foundation for Arts and Sciences (to K.K.). This work was supported by MEXT KAKENHI (No. 221S0002).

## Details of Author Contributions

Conceived and designed the experiments: RKD, KK, HM; performed the experiments: RKD, KK, RO, YO, TS, TO, TI, PS; analyzed the data RKD, KK, YO, ZB, RY, KN, TH, HM; contributed reagents/materials/analysis tools: RKD, KK, TS, TVP, EK, AH, MT, AS, HA, TTB, LTN, KVV, DQDH, TS, YY, HM; prepared the manuscript: RKD, KK, TH, HM.

